# Critical role for Piccolo in Synaptic Vesicle Retrieval

**DOI:** 10.1101/330050

**Authors:** Frauke Ackermann, Kay O. Schink, Christine Bruns, Zsuzsanna Izsvák, F. Kent Hamra, Christian Rosenmund, Craig C. Garner

## Abstract

Loss of function of the presynaptic active zone protein Piccolo has recently been linked to a devastating disease causing brain atrophy. Here, we address how Piccolo inactivation adversely affects synaptic function and thus may contributes to neuronal loss. Our analysis shows that Piccolo is critical for the activity dependent recycling and maintenance of synaptic vesicles (SVs). Specifically, we find that boutons lacking Piccolo have deficits in the Rab5/EEA1 dependent formation of early endosomes and thus the recycling of SVs. Mechanistically, impaired Rab5 function was caused by the reduced synaptic recruitment of Pra1, known to interact selectively with the zinc fingers of Piccolo. Importantly, over-expression of GTPase deficient Rab5 or the Znf1 domain of Piccolo restores the size and recycling of SV pools. These data provide a molecular link between the active zone and endosome sorting at synapses providing hints to how Piccolo contributes to both developmental and psychiatric disorders.

**Impact Statement:** The efficient recycling of synaptic vesicle proteins is critical for the integrity and reliability of synaptic transmission. Increasingly genetic and environmental insults have been shown to affect this recycling pathway, resulting in both cognitive impairment in humans and neurodegenerative diseases, yet the underlying mechanisms are poorly understood. Here we could show that the presynaptic active zone protein Piccolo regulates efficient recycling of synaptic vesicles via Pra1 and Rab5, perhaps explaining why Piccolo loss of function contributes to Pontocerebellar Hypoplasia and major depressive disorders.

## Introduction

Piccolo, also known as Aczonin, is a scaffolding protein of the cytomatrix assembled at the active zone (CAZ), a specialized region within the presynaptic terminal where SVs fusion takes place (Ackermann, Waites, & Garner, 2015; Gundelfinger, Reissner, & Garner, 2015). It is a multi-domain protein consisting of 10 Piccolo/Bassoon homology domains (PBH), two zinc-finger (Znf) domains, three coiled-coil (CC) domains, a single PDZ domain and two C2 domains (C2A/C2B) (Fenster et al., 2000; Fenster & Garner, 2002; Wang et al., 1999). Piccolo is exclusively localized to active zones (Cases-Langhoff et al., 1996; Dick et al., 2001; Hagiwara, Fukazawa, Deguchi-Tawarada, Ohtsuka, & Shigemoto, 2005; Juranek et al., 2006; Limbach et al., 2011; Nishimune, 2012a, 2012b; Siksou et al., 2007), where it forms a sandwich-like structure enclosing Bassoon (Nishimune, Badawi, Mori, & Shigemoto, 2016).

At present, the function of Piccolo is not well understood. Initial genome-wide association studies (GWAS) have linked missense mutations in Piccolo to psychiatric and developmental disorders including major depressive and bipolar disorder (Choi et al., 2011; Giniatullina et al., 2015; Minelli et al., 2012; Sullivan et al., 2009; Woudstra et al., 2013). More recently, a non-sense mutation (chr7:82579280 G>A), predicted to eliminate the C-terminal half of Piccolo, was identified in patients with Pontocerebellar Hypoplasia 3 (PCH3)(Ahmed et al., 2015). PCH3 is a devastating developmental disorder associated with profound cognitive and motor impairment, atrophy of the cerebrum and cerebellum and progressive microcephaly (Ahmed et al., 2015; Maricich et al., 2011; Rudnik-Schoneborn, Barth, & Zerres, 2014). How an active zone protein like Piccolo contributes to these diseases is still unclear.

Clues from cellular, biochemical and molecular studies indicate a structural role for Piccolo in organizing the active zone (AZ) (Ackermann et al., 2015; Gundelfinger et al., 2015) as it`s loss in the retina disrupts the assembly of photoreceptor cell ribbons (Regus-Leidig et al., 2014) and its association with other CAZ proteins, including Bassoon and CAZ-associated structural protein (CAST), (Takao-Rikitsu et al., 2004; Wang et al., 2009). However, its association with the SV priming factors RIM1 and Munc13-1 (Sudhof & Rizo, 2011) implies an additional role in SV priming, though a direct role in regulating neurotransmitter release has not been demonstrated (Mukherjee et al., 2010). Intriguingly, structure/function studies have shown that Piccolo directly interacts with a number of actin-binding proteins including Profilin-2, Daam1, Abp1 and Trio and is critical for the activity dependent assembly of presynaptic F-actin (Fenster et al., 2003; Leal-Ortiz et al., 2008; Terry-Lorenzo et al., 2016; Wagh et al., 2015; Waites, Leal-Ortiz, Andlauer, Sigrist, & Garner, 2011; Wang et al., 1999). Moreover, it was also found to interact with the prenylated Rab acceptor protein (Pra1)(Fenster et al., 2000), and the Arf GTPase-activating protein 1 (GIT1)(Kim et al., 2003). These latter features suggest roles for Piccolo in the activity dependent recycling and maintenance of SVs, which is a requirement for synapse integrity and neuronal survival.

So far, membrane trafficking within presynaptic boutons is only poorly understood. Early studies show endosomal structures at presynaptic boutons (Heuser & Reese, 1973) as well as the presence of small Rab GTPases, including Rab4, 5, 7, 10, 11, 35, 26 and 14 (Pavlos & Jahn, 2011). Members of the small Rab GTPase family are organizers of membrane trafficking tasks as they function as molecular switches alternating between a GDP bound “off-state” and a GTP bound “on-state” (Stenmark, 2009). Active, they operate together with phosphoinositides (PIs), which are also present at synapses (Uytterhoeven, Kuenen, Kasprowicz, Miskiewicz, & Verstreken, 2011; Wucherpfennig, Wilsch-Brauninger, & Gonzalez-Gaitan, 2003), to spatially and temporally recruit effector proteins like Early Endosome Antigen 1 (EEA1) (Christoforidis, McBride, Burgoyne, & Zerial, 1999; Spang, 2009) providing identity to compartments (Balla, 2013; Schink, Tan, & Stenmark, 2016; Zerial & McBride, 2001).

Surprisingly, perturbation of endosome proteins as well as over-expression of GTPase deficient Rab5 (Rab5^Q79L^) affects synapse function, causing a smaller readily releasable pool of vesicles (RRP), reduced FM uptake (Hoopmann et al., 2010; Rizzoli & Betz, 2002; Wucherpfennig et al., 2003), or an increased synaptic release probability (Wucherpfennig et al., 2003)(see also (Sasidharan et al., 2012)). These and related studies support the concept that membrane trafficking is a fundamental feature of presynaptic boutons, helping them maintain functional fidelity over time.

To better understand the synaptic function of Piccolo and its contribution to various psychiatric diseases as well as PCH3, we recently generated line of rats in which the *Pclo* gene was disrupted by transposon mutagenesis (*Pclo*^*gt/gt*^)(Medrano et al., 2018). Our initial analysis of these animals reveals that they exhibit many of the hallmarks also seen in PCH3 patients including microcephaly, seizures and reduced volume of the cerebrum, cerebellum and pons (Falck, Ackermann & Garner in prep), suggesting this animal may be a good model of PCH3. In the current study, we used these animals to gain deeper insights into the molecular mechanisms of how Piccolo loss of function alters synaptic function and perhaps contributes to neurodevelopmental and psychiatric disorders.

Our investigations reveal that knockout of Piccolo causes a dramatic decrease in the number of SVs per synaptic bouton as well as the size of the total recycling pool (TRP) of SVs. Mechanistically, we found that these phenotypes are caused by the reduced recruitment of the Piccolo binding partner Pra1 to boutons, the activation of Rab5 as well as the formation of early endosome. These data provide a molecular link between the active zone protein Piccolo, endosome trafficking and the functional maintenance of SV pools providing a possible molecular mechanism for how Piccolo loss could contribute to PCH3 and major depressive disorders among others.

## Results

### Generation and characterization of the Piccolo Knockout rat

Transposon mutagenesis was recently used to generate rats with a disrupted *Pclo* gene (*Pclo*^*gt/gt*^)(Medrano et al., 2018). As shown in Fig. 1 A, the transposon element became integrated into exon 3 of the *Pclo* genomic sequence, leading to a stop in the reading frame. The genotyping of offspring by PCR from genomic DNA led to the amplification of a single 280 bp band from *Pclo*^*wt/wt*^ rats, two bands of 280 bp and 450 bp from heterozygous *Pclo*^*wt/gt*^ animals and a single 450 bp band from *Pclo*^*gt/gt*^ animals (Fig. 1 B). Western blot analysis from postnatal day 2 (P2) rat brain as well as primary hippocampal neuron lysates confirms the loss of Piccolo full-length protein. As reported earlier (Waites et al., 2011), prominent bands at 560 kD as well as other bands between 70-450 kD were detected in lysates from *Pclo*^*wt/wt*^ brains (Fig. 1 C, D), reflecting the expression of multiple Piccolo isoforms from the *Pclo* gene (Fenster & Garner, 2002). A similar banding pattern, though less intense, was seen in brain lysates from *Pclo*^*wt/gt*^ animals (Fig. 1 C). In brain lysates from *Pclo*^*gt/gt*^ animals, the dominant bands at 560 and 450 kD were absent, indicating the loss of the predominant Piccolo isoforms, though a few weaker bands at 300, 90 and 70 kD are still present (Fig. 1 C). However, the loss of Piccolo from *Pclo*^*gt/gt*^ brains and hippocampal neurons (Fig. 1, C and D) does not appear to have an impact on the gross anatomy of the hippocampus (Fig. S1 A and B). Moreover, immuno-staining of hippocampal neurons with antibodies against the central region of Piccolo, the microtubule-associated protein 2 (MAP2) and the vesicular glutamate transporter 1 (VGlut1) revealed the presence of Piccolo immuno-reactive puncta along the dendrites of *Pclo*^*wt/wt*^ neurons that co-localized with the presynaptic vesicle protein VGlut1 (Fig. 1 E). In *Pclo*^*gt/gt*^ neurons, the staining intensity of Piccolo co-localizing with VGlut1 puncta was reduced by more than 70% (Fig. 1 E), confirming the loss of most Piccolo isoforms from these synapses. Similarly, Piccolo immuno-reactivity is gone from hippocampal brain section (Fig. S1, C and D).

**Figure 1.**
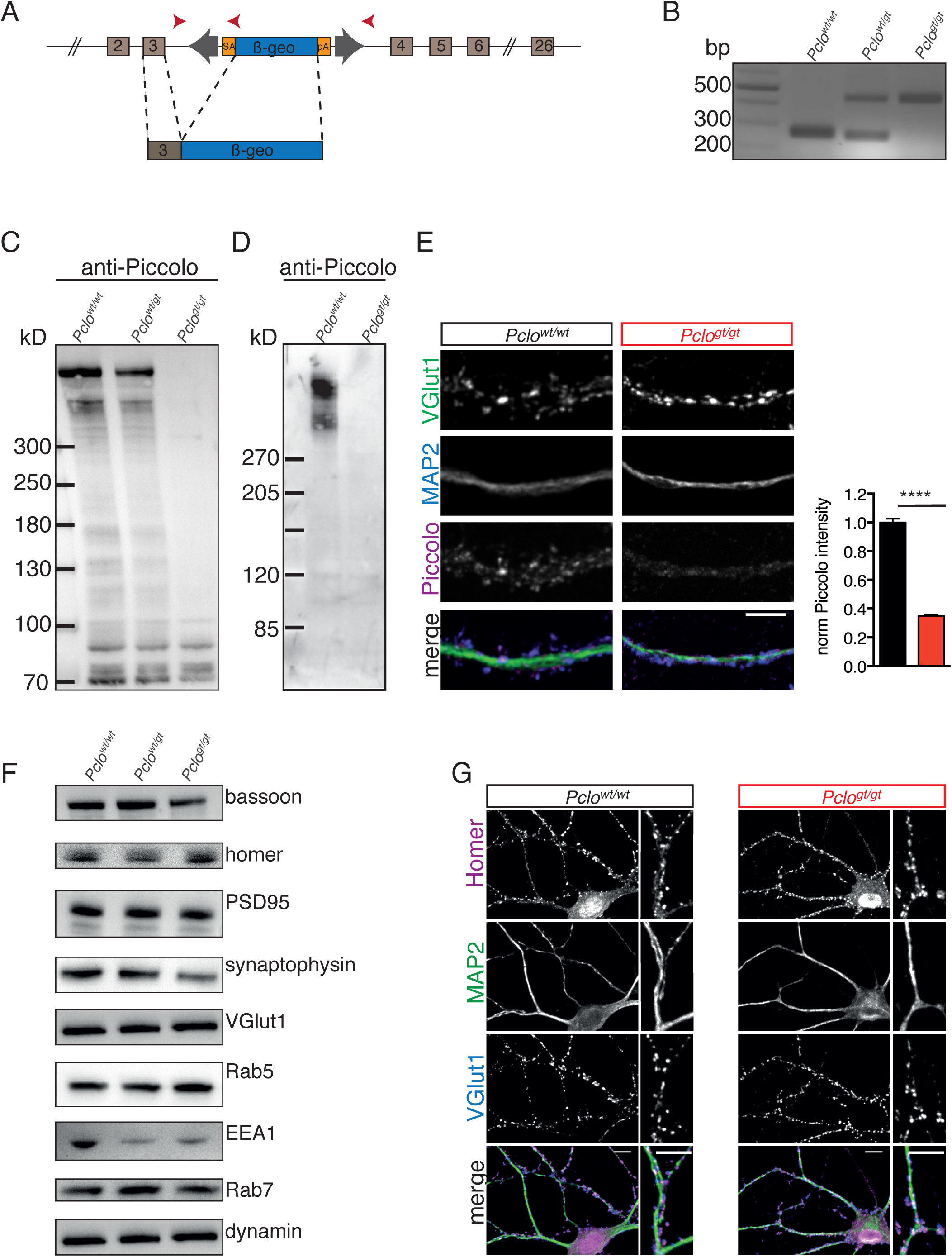
Transposon mutagenesis leads to the loss of Piccolo protein in rats. (A) Transposon insertion into exon 3 of *Pclo* genomic sequence causing a stop codon. (B) PCR verification of genotypes. PCR from *Pclo*^*wt/wt*^ *Pclo*^*wt/gt*^ and *Pclo*^*gt/gt*^ genomic DNA results in the amplification of a 260 bp WT and/or a 450 bp KO specific bands, respectively. (C) Western blot analysis of P0-P2 brain lysates. Bands corresponding to Piccolo isoforms (70-560 kDa) are detectable in *Pclo*^*wt/wt*^ and *Pclo*^*wt/gt*^ lysates but not in *Pclo*^*gt/gt*^ lysates (n = 3 independent experiments). (D) Western blot analysis of neurons lysates. Bands corresponding Piccolo isoforms (560 - 300 kDa) are detectable in *Pclo*^*wt/wt*^ but not in *Pclo*^*gt/gt*^ lysates (n = 3 independent experiments). (E) Immunocytochemical stainings of hippocampal neurons. Piccolo co-localizes with VGlut1 in *Pclo*^*wt/wt*^ neurons but not in *Pclo*^*gt/gt*^ neurons (right panel). Left panel, quantification of normalized Piccolo intensities at VGlut1 puncta (*Pclo*^*wt/wt*^ = 1 ± 0.03, n = 586 synapses; *Pclo*^*gt/gt*^ = 0.35 ± 0.01, n = 718 synapses; p < 0.0001). (F) Western blot analysis of P0-P2 brain lysates. Expression levels of Homer, PSD95, VGlut1, Dynamin and Rab7 are not altered, whereas the levels of Synaptophysin and EEA1 are decreased and Rab5 levels are increased in *Pclo*^*gt/gt*^ brain lysates (n = 3 independent experiments). (G) Immunocytochemical stainings of hippocampal neurons. The density of Homer and VGlut1 opposing puncta along dendrites (MAP2) is comparable between *Pclo*^*wt/wt*^ and *Pclo*^*gt/gt*^ neurons. Scale bar in E and G 10 μm, scale bar in G (zoom window) 5 μm. Mean ± SEM, Students T-test.

As a core protein of the AZ, we were keen to examine the effect of loss of Piccolo on the expression levels of other synaptic proteins. In Western blot analysis of P2 rat brain lysates, most synaptic proteins tested, including Homer, PSD95, Dynamin as well as VGlut1 were not affected (Fig. 1 F). Intriguingly, Piccolo deficiency was associated with a decreased expression of Bassoon, the synaptic vesicle protein Synaptophysin and the early endosome marker EEA1, whereas the levels of the small GTPase Rab5 were increased (Fig. 1 F). As an initial indication of whether these changes affected synapse formation, hippocampal cultures were immuno-stained with MAP2, VGlut1 and the postsynaptic marker Homer. VGlut1 puncta opposing Homer puncta were present in *Pclo*^*wt/wt*^ and *Pclo*^*gt/gt*^ neurons (Fig. 1 G), indicating that most Piccolo isoforms are not required for the initial assembly of excitatory synapses. This is further supported by the fact, that within CA1 and CA3 of the hippocampus VGlut1 staining is not overtly changed (Fig. S1, C and D).

### Ultrastructural analysis of Pclo^gt/gt^ synapses

To more directly assess a role of Piccolo in synapse assembly, we analyzed the ultrastructure of synapses by EM using high pressure freezing and freeze substitution. EM micrographs from *Pclo*^*wt/wt*^ and *Pclo*^*gt/gt*^ neurons reveal the presence of synaptic junctions, with prominent postsynaptic densities (PSDs) formed onto axonal varicosities of *Pclo*^*gt/gt*^ neurons, with similar dimensions (length) to *Pclo*^*wt/wt*^ synapses (Fig. 2 A, B, E). At many *Pclo*^*gt/gt*^ synapses, synaptic vesicles (SVs) (< 50nm diameter) are detected (Fig. 2 B), however the number of SVs/bouton is significantly reduced (Fig. 2, B and C). An analysis of the number of SVs docked at AZs revealed that the loss of Piccolo does not affect the number of readily releasable vesicles (Fig. 2 D, H, I). Intriguingly, lower SV density in *Pclo*^*gt/gt*^ is accompanied by a high number of endosome like structures with a diameter larger than 60 nm (Fig. 2 B, F, G). These data suggest that Piccolo is required for the maintenance and/or recycling of SVs.

**Figure 2.**
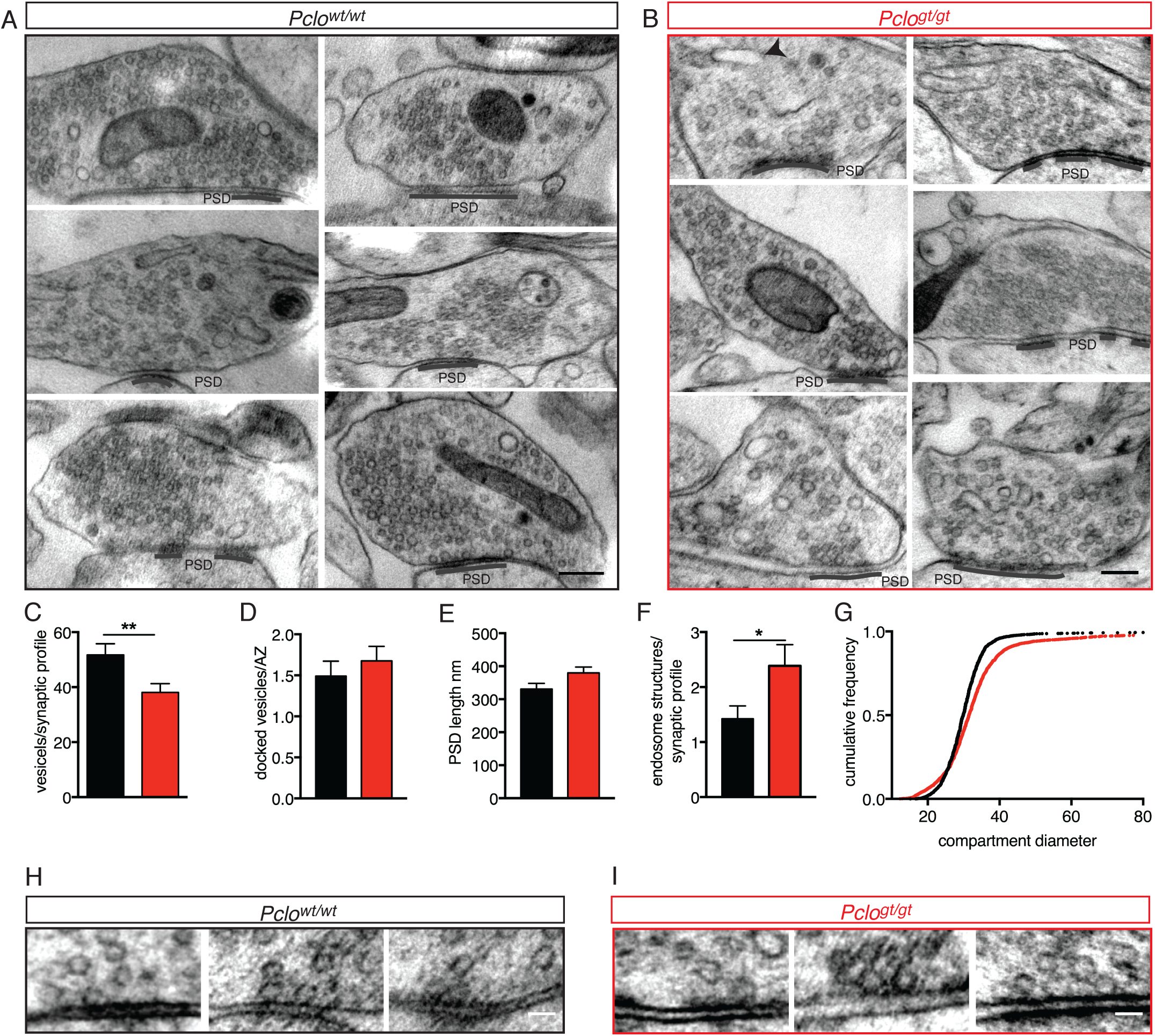
*Pclo*^*gt/gt*^ synapses display morphological changes on ultrastructural level. (A,B) Example electron micrographs of *Pclo*^*wt/wt*^ (A) and *Pclo*^*gt/gt*^ (B) synapses. (C-G) Quantification of electron micrographs. (C) Synaptic vesicle density is decreased in *Pclo*^*gt/gt*^ terminals (*Pclo*^*wt/wt*^ = 51.73 ± 4.05, n = 52 synapses; *Pclo*^*gt/gt*^ = 38.08 ± 3.19, n = 59 synapses; p = 0.0085). (D) The number of docked vesicles per active zone is not altered (*Pclo*^*wt/wt*^ = 1.49 ± 0.18, n = 59 AZs; *Pclo*^*gt/gt*^ = 1.67 ± 0.18, n = 40 AZs). (E) The total length of the PSDs is not altered (*Pclo*^*wt/wt*^ = 331 ± 17.09 nm, n = 72 PSDs; *Pclo*^*gt/gt*^ = 379.6 ± 18.22 nm, n = 69 PSDs). (F) The number of endosome structures is increased at Piccolo *Pclo*^*gt/gt*^ synapses (*Pclo*^*wt/wt*^ = 1.42 ± 0.24, n = 52 synapses; *Pclo*^*gt/gt*^ = 2.39 ± 0.38, n = 57 synapses). (G) Cumulative blot depicting the distribution of endosome compartment diameters. *Pclo*^*gt/gt*^ vesicular compartments show a shift towards larger diameters (*Pclo*^*wt/wt*^: n = 2729 compartments measures; *Pclo*^*gt/gt*^: n = 2387 compartments measured). (H, I) Example electron micrographs showing docked vesicles at *Pclo*^*wt/wt*^ (H) and *Pclo*^*gt/gt*^ AZs (I). Scale bar in A and B 200 nm, scale bar in D and E 50 nm. Mean ± SEM, Students *t*-test.

### Pools of recycling SVs are reduced but synaptic release properties are unchanged in boutons lacking Piccolo

Given its restricted and tight association with presynaptic AZs (Cases-Langhoff et al., 1996), we also explored whether loss of Piccolo adversely affects synaptic transmission. To this end, we performed whole cell patch clamp recordings from cultures of autaptic hippocampal neurons. Under these conditions, we observed a slight, but non-significant, reduction in excitatory postsynaptic currents (EPSC) amplitudes elicited by a single action potential (AP) in *Pclo*^*gt/gt*^ compared to *Pclo*^*wt/wt*^ neurons (Fig. 3 A). The application of 5 s pulses of hypertonic sucrose (500mM, ∼800 mOsm) to determine the size of the readily releasable pool (RRP) of SVs revealed that it was not altered in *Pclo*^*gt/gt*^ neurons (Fig. 3 D), consistent with the unaltered umber of docked vesicles per active zone (Fig. 2D). Similarly, SV release probability (Pvr), as a measure for release efficiency, is not changed in neurons generated from *Pclo*^*gt/gt*^ animals (Fig. 3 C). In addition, the paired pulse ratio (PPR) (25 ms pulse interval) was also not significantly altered (Fig. 3 B). However, the steady state EPSC amplitude at the end of a 10-Hz/5s train stimulation in *Pclo*^*gt/gt*^ compared to *Pclo*^*wt/wt*^ neurons was reduced, indicating impaired maintenance of synaptic transmission during intense vesicle release (Fig. 3 E). Together, these data indicate that the release properties of SVs are not severely changed at AZs lacking Piccolo, though the sustained release of neurotransmitter is compromised, perhaps by the reduced number of SVs/bouton (Fig. 2, B and C) or their efficient recycling.

**Figure 3.**
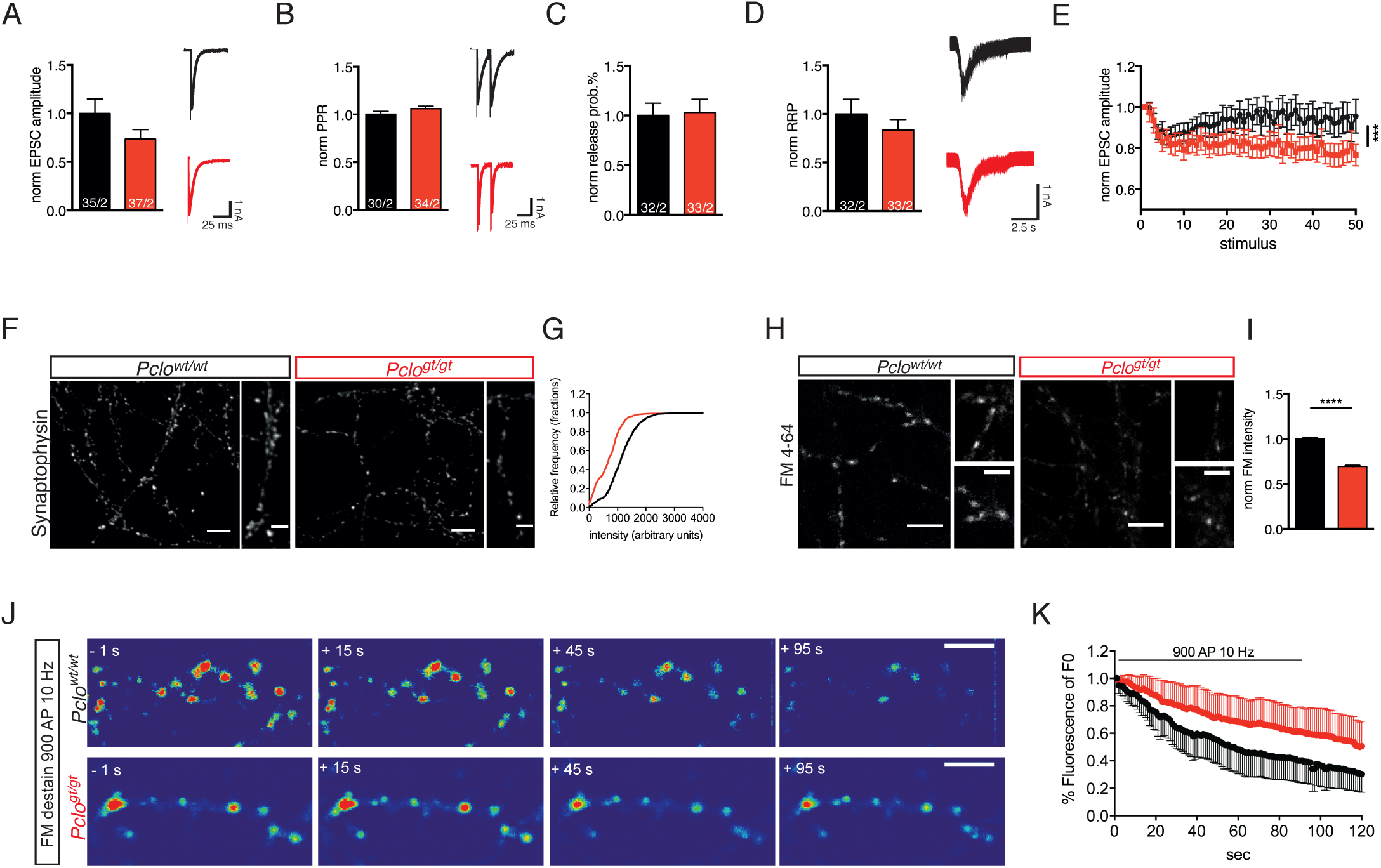
Synaptic vesicle release properties are not altered, but the total recycling pool of vesicles is reduced due to the loss of Piccolo. (A-E) Patch clamp recordings on autaptic primary hippocampal neurons. (A) Amplitudes of evoked postsynaptic currents (EPSCs) are reduced in *Pclo*^*gt/gt*^ neurons (*Pclo*^*wt/wt*^ = 1.0 ± 0.15, *Pclo*^*gt/gt*^ = 0.73 ± 0.10; p = 0.14, 2 independent experiments). (B) Paired pulse ratio is not altered in *Pclo*^*gt/gt*^ neurons (*Pclo*^*wt/wt*^ = 1.00 ± 0.03, *Pclo*^*gt/gt*^ = 1.06 ± 0.03; p = 0.15, 2 independent experiments).(C) Vesicle release probability is not changed upon Piccolo loss (*Pclo*^*wt/wt*^ = 1± 0.13, *Pclo*^*gt/gt*^ = 1.03 ± 0.13; p = 0.87, 2 independent experiments). (D) The readily releasable pool of vesicles (RRP) in *Pclo*^*gt/gt*^ neurons is not altered (*Pclo*^*wt/wt*^ = 1 ± 0.15, *Pclo*^*gt/gt*^ = 0.83 ± 0.11; p = 0.37, 2 independent experiments). (E) Loss of Piccolo causes a increase in EPSC amplitude depression during 10 Hz train stimulation (*Pclo*^*wt/wt*^ 34 cells, *Pclo*^*gt/gt*^ 35 cells; 2 independent experiments). (F) Immunocytochemical stainings of hippocampal neurons. (G) The cumulative distribution of Synaptophysin intensities is shifted towards lower intensities in *Pclo*^*gt/gt*^ neurons (*Pclo*^*wt/wt*^: 1341 puncta; *Pclo*^*gt/gt*^: 712 puncta; 4 independent experiments). (H) FM 4-64 dye uptake experiments. (I) Quantification of (H). Dye uptake is significantly reduced in *Pclo*^*gt/gt*^ neurons (*Pclo*^*wt/wt*^ = 1 ± 0.01, n = 1026 puncta; *Pclo*^*gt/gt*^ = 0.69 ± 0.01, n = 867 puncta; p < 0.0001; 4 independent experiments). (J) Selected images of synaptic boutons releasing loaded FM1-43 dye during a 900 AP 10 Hz stimulation. (K) Quantification of changes in FM1-43 dye intensities per bouton over time. Note, FM1-43 destaining rate from boutons is slower in *Pclo*^*gt/gt*^ versus *Pclo*^*wt/wt*^ neurons. In *Pclo*^*wt/wt*^ neurons, about 70% of the initially loaded FM1-43 dye is released 120 seconds after start of the stimulation (*Pclo*^*wt/wt*^ = 0.30 ± 0.13, n = 402 synapses, 3 independent experiments), whereas in *Pclo*^*gt/gt*^ neurons less than 30% is released (*Pclo*^*gt/gt*^ = 0.51 ± 0.18, n = 215 synapses, 3 independent experiments). Scale bar in F, H and J 10 μm, scale bar in zoom in F and H 5 μm. Numbers in bar graphs represent number of cells/number of cultures. Mean ± SEM, Student`s *t* -test.

To explore whether reduced pools and/or the recycling of SVs contribute to the observed changes in boutons lacking Piccolo, we performed two types of experiments. Initially, immunocytochemistry was used to measure the total pool of SVs by staining cultures with antibodies against the SV protein Synaptophysin. This revealed a dramatic (48 %) reduction of Synaptophysin content per bouton (Fig. 3, F and G), indicating that SV loss (as seen in EM micrographs) (Fig. 2, A and B) is a putative contributor to the reduced functionality of *Pclo*^*gt/gt*^ neurons. Next, we performed FM-dye uptake experiments (Smith & Betz, 1996) to determine the size of the total recycling pool (TRP) of SVs. Here, *Pclo*^*gt/gt*^ neurons display a 30% reduction in the FM4-64 intensity, consistent with a smaller TRP of SVs (Fig. 3, H and I). Finally, to assess whether SVs recovered during endocytosis are efficiently recycled back to the SV cluster, we also determined the destaining kinetics of boutons, loaded with FM1-43, during 10Hz/900AP field stimulation. Remarkably, boutons lacking Piccolo released the FM dye at a much slower rate than WT boutons (Fig. 3 J, K). As SV exocytosis is normal in boutons lacking Piccolo, these data indicate that the recycling of SV must be compromised, as dye loaded into boutons is not efficiently released.

### Levels of endosome proteins are reduced in *Pclo*^*gt/gt*^ synapses

Important questions raised by our initial analysis are a) why SV pool size is smaller in boutons lacking Piccolo and b) why FM unloading rates are slowed? Based on the increased presence of endosome-like membranes (Fig. 2 B), one potential explanation could be a defect in the recycling or reformation of SVs. As an initial test of this hypothesis, we expressed Rab5, tagged with GFP, to monitor the presence of endocytic compartments (Spang, 2009; Stenmark, 2009) in *Pclo*^*gt/gt*^ neurons. Interestingly, we observed an increase in the number of GFP-Rab5 positive puncta along axons of *Pclo*^*gt/gt*^ versus *Pclo*^*wt/wt*^ neurons (Fig. 4, A and B). To examine whether this was associated with a general increase in the endo-lysosomal trafficking of SVs, we monitored the presence of GFP-Rab7, a marker for late endosomes (Zerial & McBride, 2001)(Fig. 4 B). Surprisingly, the number of GFP-Rab7 puncta along axons was reduced in *Pclo*^*gt/gt*^ neurons (Fig. 4, A and B). These data indicate that the maturation of membranes within the endosome compartment from Rab5 positive endosomes towards late Rab7 positive endosomes is affected by the loss of Piccolo.

**Figure 4.**
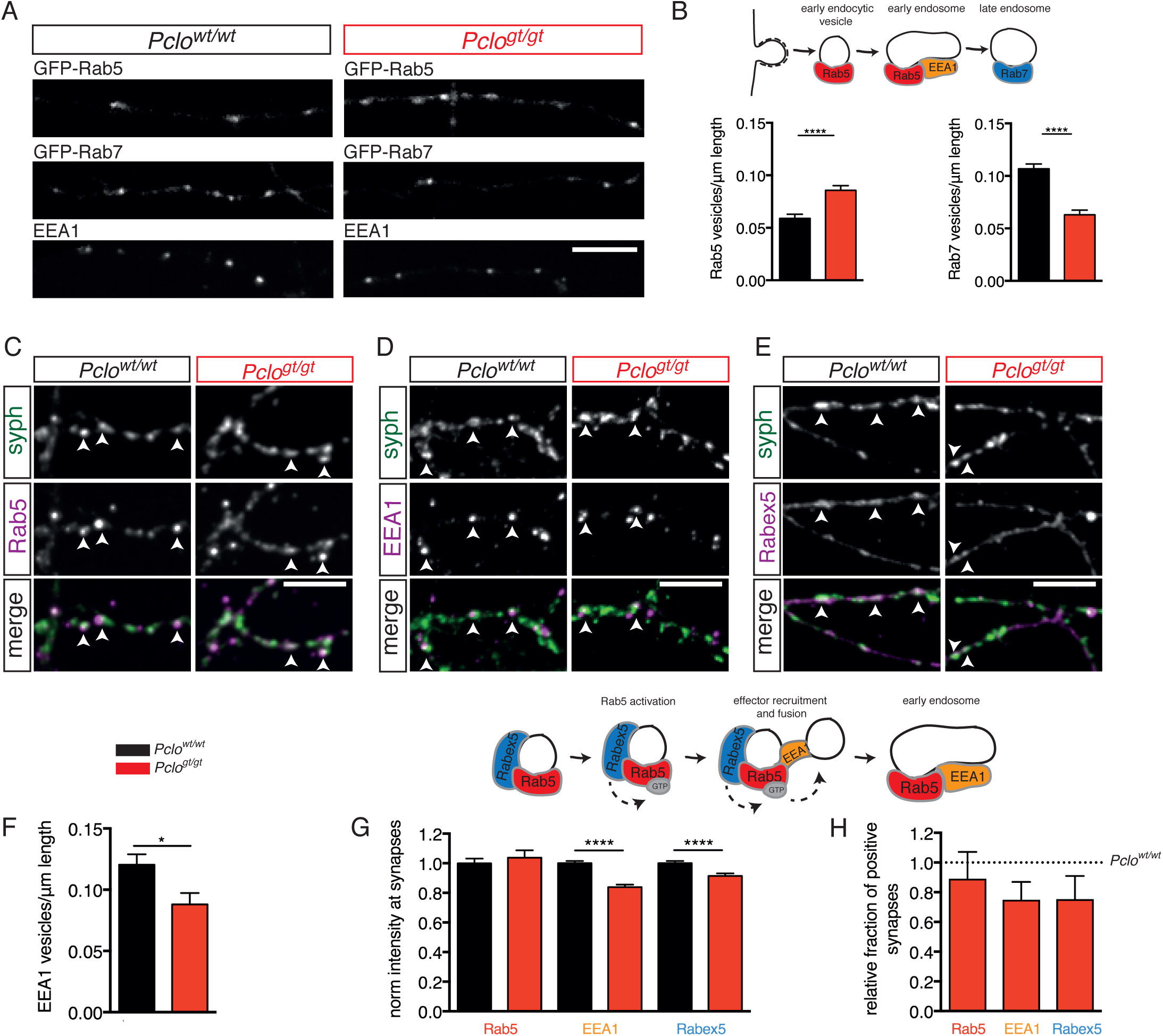
Levels of endosome proteins are reduced at synapses lacking Piccolo. (A)Images of axon segments from hippocampal neuron expressing GFP-Rab5 or GFP-Rab7 or immune-stained with antibodies to EEA1. (B)Schematic of SV membrane trafficking and quantification of data in A. Upper panel, schematic illustrates when Rab5, EEA1 and Rab7 become associated with endosomal membranes. Lower panels, in *Pclo*^*wt/wt*^ neurons less GFP-Rab5 puncta are present per unit length of axon (*Pclo*^*wt/wt*^ = 0.06 ± 0.004, n = 139 axon sections; *Pclo*^*gt/gt*^ = 0.08 ± 0.004, n = 143 axon sections; p < 0.0001; 3 independent experiments). In *Pclo*^*wt/wt*^ neurons more GFP-Rab7 puncta are present per unit length of axon (*Pclo*^*wt/wt*^ = 0.11 ± 0.004, n= 139 axon sections; *Pclo*^*gt/gt*^ = 0.06 ± 0.004, n = 142 axon sections; p < 0.0001; 2 independent experiments). (C-E) Immunocytochemical staining of hippocampal neurons for synaptic and endosome proteins. (C) Rab5 is present at *Pclo*^*wt/wt*^ and *Pclo*^*gt/gt*^ synapses, no difference in their intensities is detectable. (D) EEA1 intensities are significantly reduced in *Pclo*^*gt/gt*^ synapses.(E) Rabex5 intensities are significantly reduced in *Pclo*^*gt/gt*^ synapses. (F) Quantification of A. In *Pclo*^*wt/wt*^ neurons more EEA1 positive puncta are present per unit length of axon compared to *Pclo*^*gt/gt*^ neurons (*Pclo*^*wt/wt*^ = 0.12 ± 0.01, n = 29 axon sections; *Pclo*^*gt/gt*^ = 0.09 ± 0.004, n = 34 axon sections; p = 0.01; 2 independent experiments). (G) Schematic of Rab5 complexes during membrane trafficking upper panel, and quantification of (C, D and E) lower panel. The levels of Rab5 are not altered between *Pclo*^*wt/wt*^ and *Pclo*^*gt/gt*^ synapses (*Pclo*^*wt/wt*^ = 1 ± 0.03, n = 808 synapses; *Pclo*^*gt/gt*^ = 1.04 ± 0.05, n = 773 synapses; p = 0.5242; 3 independent experiments). EEA1 intensity is significantly reduced in *Pclo*^*gt/gt*^ synapses (*Pclo*^*wt/wt*^ = 1 ± 0.01, n = 4371 synapses; *Pclo*^*gt/gt*^ = 0.84 ± 0.02, n = 2960 synapses; p < 0.0001; 7 independent experiments). Rabex5 intensity is reduced in *Pclo*^*gt/gt*^ synapses (*Pclo*^*wt/wt*^ = 1 ± 0.01, n = 2397 synapses; *Pclo*^*gt/gt*^ = 0.91 ± 0.02, n = 2551 synapses; p = 0.0002; 7 independent experiments). (H) The fraction of synapses positive for Rab5, EEA1 and Rabex5 is reduced in neurons lacking Piccolo. Relative fraction of Rab5 positive synapses: 0.89 ± 0.18, p = 0.5032, n = 3 independent experiments. Relative fraction of EEA1 positive synapses:0.75 ± 0.12, p = 0.1517, n = 7 independent experiments. Relative fraction of Rabex5 positive synapses is: 0.75 ± 0.16, p = 0.2744, n = 7 independent experiments. Scale bar represents 10 μm. Mean ± SEM, Student`s *t* -test.

To gain insights into the possible mechanism, we examined EEA1, a docking factor that facilitates the homotypic fusion between early endocytic vesicles, which then become early endosomes (Christoforidis et al., 1999). We quantified the abundance of EEA1 puncta in *Pclo*^*gt/gt*^ neurons (Fig. 4 A, F). Indeed, the number of EEA1 immuno-positive puncta along axons was also reduced in *Pclo*^*gt/gt*^ neurons (Fig. 4, A and F), further indicating that the loss of Piccolo alters endocytic membrane trafficking.

As Piccolo is an AZ protein, we next analyzed whether the levels of Rab5 and EEA1 were also changed at synapses. Here, *Pclo*^*wt/wt*^ and *Pclo*^*gt/gt*^ neurons were immuno-stained with antibodies against endogenous Rab5 and EEA1 as well as Synaptophysin. Quantifying intensities at Synaptophysin spots revealed that while Rab5 levels were not altered (Fig. 4, C and G), EEA1 levels were decreased in *Pclo*^*gt/gt*^ versus *Pclo*^*wt/wt*^ synapses (Fig. 4, D and G). Note, though the fraction of synapses positive for Rab5 was only modestly affected in *Pclo*^*gt/gt*^ neurons, those positive for EEA1 were decreased by more than 20% compared to *Pclo*^*wt/wt*^ neurons (Fig. 4 H).

The recruitment of EEA1 to endocytic vesicles is dependent on GTP bound active Rab5 (Mishra, Eathiraj, Corvera, & Lambright, 2010; Murray et al., 2016; Simonsen et al., 1998). We thus examined whether the levels of the GEF known to regulate Rab5 activity, Rabex5 (Horiuchi et al., 1997), were also changed in synapses of *Pclo*^*gt/gt*^ neurons. Here, we observed that the levels, as well as the fraction of Rabex5 positive synapses, were decreased (Fig. 4 E, G, H). Taken together these data indicate that Piccolo normally contributes to the efficient recycling of endosome membranes within presynaptic boutons. Consistent with this concept, no differences in the levels of Rab5, EEA1 or Rabex5 were detected in the cell soma and along dendrites between *Pclo*^*wt/wt*^ and *Pclo*^*gt/gt*^ neurons (Fig. S2).

### PI(3)P positive endosome organelles are smaller at Piccolo KO synapses

PI(3)P is one of the earliest marker of early endosomes (Schink, Raiborg, & Stenmark, 2013). It can be readily detected through the recombinant expression of the PI(3)P binding domain (FYVE) of the ESCRT protein hepatocyte growth factor-regulated tyrosine kinase substrate (Hrs) (Komada & Soriano, 1999), making it a specific tool to visualize early endosomes in cells (Fig. S3 A). To examine whether the reduction of EEA1 levels could be due to abnormally low levels of PI(3)P in synaptic boutons lacking Piccolo, GFP-2x-FYVE was expressed in primary hippocampal neurons, where it labels vesicles along axons (Fig. S3 B). No differences in the number of GFP-2x-FYVE organelles per axon length could be detected between *Pclo*^*wt/wt*^ and *Pclo*^*gt/gt*^ neurons (Fig. S3, B and C), suggesting that PI(3)P positive organelles form normally in *Pclo*^*gt/gt*^ neurons.

However, to gain further insight into the size and distribution of PI(3)P organelles at synapses, we performed super resolution microscopy on neurons expressing GFP-2x-FYVE. Using structure illumination microscopy (SIM), we observed that on average *Pclo*^*gt/gt*^ boutons (Bassoon and Synaptophysin positive) had significantly smaller GFP-2x-FYVE organelles (< 55 μm^2^) than *Pclo*^*wt/wt*^ boutons (up to 80 μm^2^)(Fig. S3, D and E). These data indicate an altered maturation of early endosomes in *Pclo*^*gt/gt*^ boutons, a concept consistent with the prevalence of small endocytic vesicles (60-100 nm) in electron micrographs of *Pclo*^*gt/gt*^ boutons (Fig. 2).

### Fewer endosome proteins are recruited to PI(3)P positive organelles along axons in Pclo^gt/gt^ neurons

A possible explanation for smaller PI(3)P organelles in *Pclo*^*gt/gt*^ boutons is that the transition between small early endocytic vesicles and larger early endosomes is attenuated. To test this hypothesis, we analyzed the recruitment of two endogenous endosome markers, Rab5 and EEA1, onto GFP-2x-FYVE membranes. Here, we analyzed the fraction of FYVE/Rab5 double positive organelles as a measure for early endocytic vesicles, and the fraction of FYVE/Rab5/EEA1 triple positive organelles as a measure for early endosomes (Fig. 5 A). This analysis was performed within axons, as the small size of synapses does not allow individual vesicles to be resolved. In *Pclo*^*gt/gt*^ axons, the amount of endogenous EEA1 as well as Rab5 at GFP-2x-FYVE organelles is significantly decreased (60 and 20% respectively), although GFP-2x-FYVE intensity is slightly increased (Fig. 5, B and D). Of note, although the intensity of Rab5 at PI(3)P sites is decreased in *Pclo*^*gt/gt*^ neurons, the overall fraction of endocytic vesicles that are double positive for GFP-2x-FYVE and Rab5 is not significantly altered in comparison to *Pclo*^*wt/wt*^ neurons (Fig. 5 E). In contrast, the fraction of early endosomes is reduced by about 70% in *Pclo*^*gt/gt*^ neurons (Fig. 5 F). Taken together, these data indicate that the recruitment of EEA1 towards PI(3)P sites is affected by the loss of Piccolo, slowing the maturation of early endocytic vesicles into early endosomes.

**Figure 5.**
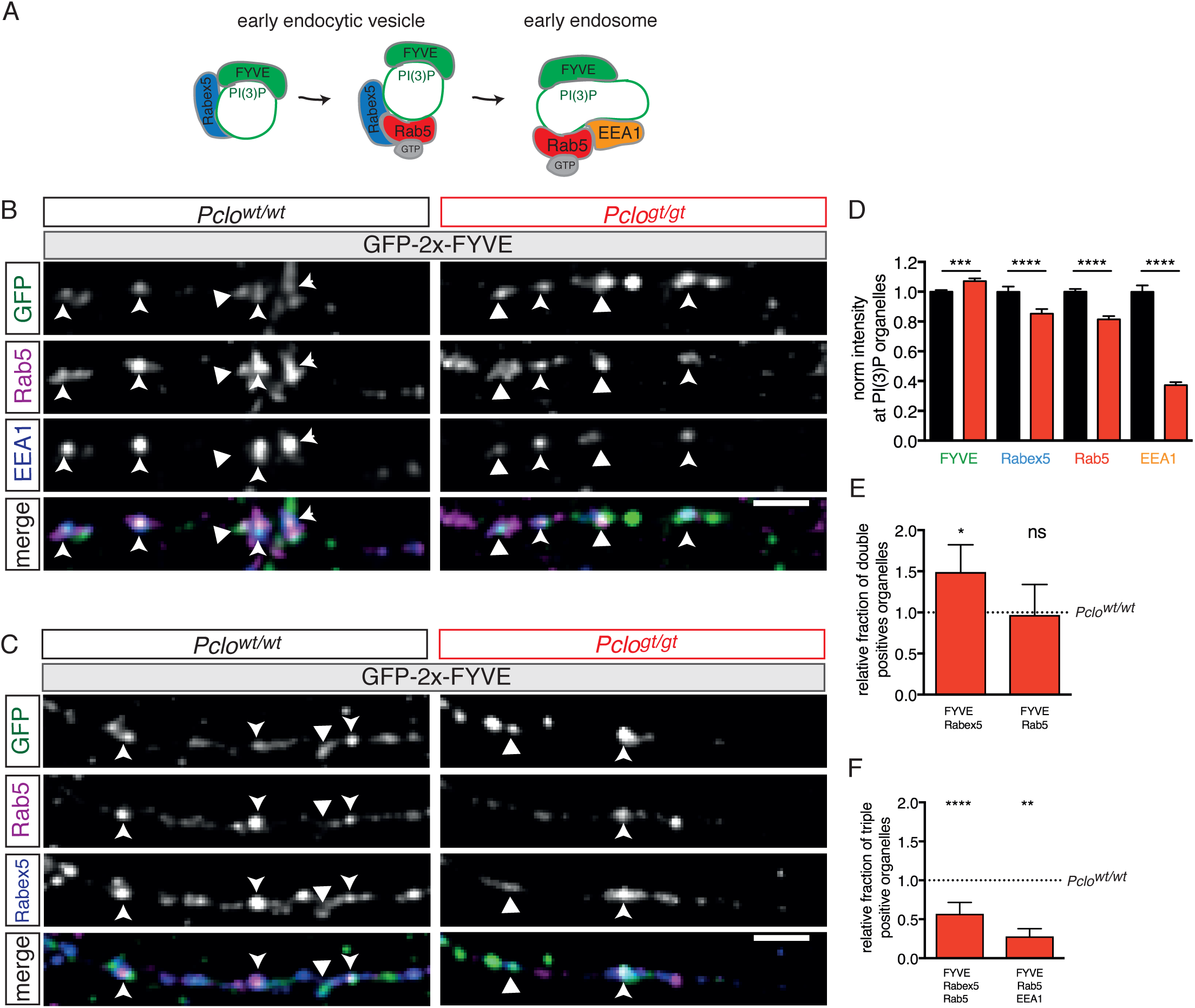
Fewer endosome proteins are recruited towards PI(3)P organelles along axons lacking Piccolo. (A) Schematic of early endocytic trafficking steps. After pinching off from the plasma membrane early endocytic vesicles undergo consecutive maturation steps. The lipid PI(3)P is generated and a stable complex consisting of Rab5 and its GEF Rabex5 is formed creating a pool of active Rab5. This step is necessary to recruit EEA1 and form early endosomes. (B) Rab5 and EEA1 intensities at GFP-2x-FYVE organelles along axons in *Pclo*^*gt/gt*^ vs *Pclo*^*wt/wt*^ neurons. (C) Rab5 and Rabex5 intensities were measured at PI(3)P organelles. (D) Quantification of B and C. The intensity of GFP-2x-FYVE is increased in *Pclo*^*gt/gt*^ neurons (*Pclo*^*wt/wt*^ = 1 ± 0.01, n = 1530 puncta; *Pclo*^*gt/gt*^ = 1.07 ± 0.02, n = 1247 puncta; p = 0.0006; 6 independent experiments). Less Rabex5 is present at PI(3)P membranes in *Pclo*^*gt/gt*^ neurons (*Pclo*^*wt/wt*^ = 1 ± 0.03, n = 1530 puncta; *Pclo*^*gt/gt*^ = 0.85 ± 0.03, n = 1247 puncta; p = 0.0017; 6 independent experiments). The levels of Rab5 at PI(3)P organelles are decreased (*Pclo*^*wt/wt*^ = 1 ± 0.02, n = 1530 puncta; *Pclo*^*gt/gt*^ = 0.81 ± 0.02, n = 1242 puncta; p < 0.0001; 6 independent experiments). The amount of EEA1 at endosome membranes is reduced (*Pclo*^*wt/wt*^ = 1 ± 0.04, n = 1237 puncta; *Pclo*^*gt/gt*^ = 0.37 ± 0.02, n = 936 puncta; p < 0.0001; 6 independent experiments). (E) Quantification of double positive compartments along axons. The fraction of GFP-2x-FYVE/Rab5 is not altered (*Pclo*^*gt/gt*^ = 0.97 ± 0.33, p = 0.72, n = 5 independent experiments). The fraction of GFP-2x-FYVE/Rabex5 double positive vesicles is increased in *Pclo*^*gt/gt*^ neurons (*Pclo*^*gt/gt*^ = 1.49 ± 0.33, p = 0.0476, n = 5 independent experiments). (F) The relative percentage of GFP-2x-FYVE/Rabex5/Rab5 as well as GFP-2x-FYVE/Rab5/EEA1 positive vesicles is decreased in *Pclo*^*gt/gt*^ neurons (GFP-2x-FYVE/Rabex5/Rab5: *Pclo*^*gt/gt*^ = 0.57 ± 0.14, p = 0.0014, n = 5 independent experiments; GFP-2x-FYVE/Rab5/EEA1: *Pclo*^*gt/gt*^ = 0.28 ± 0.10, p < 0.001, n = 6 independent experiments). Scale bars represent 10 μm. Mean ± SEM, Student`s *t* -test and ANOVA with Tukey multi comparison test.

As EEA1 recruitment to early endocytic structures depends on both PI(3)P and activated Rab5 (Mishra et al., 2010; Murray et al., 2016; Simonsen et al., 1998) and PI(3)P levels are only slightly increased in *Pclo*^*gt/gt*^ neurons (Fig. 5 D), it remains possible that Rab5 activation/activity is decreased, causing a less efficient enrichment of EEA1 towards PI(3)P sites. To test this hypothesis, we examined the levels of a known Rab5 GEF Rabex5 (Horiuchi et al., 1997) at PI(3)P sites. Similar to Rab5 and EEA1, the amounts of Rabex5 are decreased in *Pclo*^*gt/gt*^ neurons (Fig. 5, C and D). Notably, among the GFP-2x-FYVE positive organelles, those that are Rabex5 double positive are significantly increased in *Pclo*^*gt/gt*^ neurons (Fig. 5 E). In contrast, those that are triple positive for GFP-2x-FYVE, Rabex5 and Rab5 are reduced by 40% (Fig. 5 F). Together, this suggests that the reduced recruitment of EEA1 to Rab5/PI(3)P early endocytic vesicles could be due to an altered complex formation between Rab5 and its GEF, Rabex5.

### GDP-locked Rab5 (Rab5^S34N^) expression in *Pclo*^*wt/wt*^ neurons decreases early endosomes to similar levels seen in *Pclo*^*gt/gt*^ neurons

To probe the hypothesis that reduced early endosome numbers in *Pclo*^*gt/gt*^ neurons are the result of less active Rab5, we expressed a GDP-locked Rab5 (Rab5^S34N^) in *Pclo*^*wt/wt*^ neurons analyzing whether it can mimic the *Pclo*^*gt/gt*^ endosome phenotype. Interestingly, we observe that the expression of Rab5^S34N^ in *Pclo*^*wt/wt*^ neurons decreases Synaptophysin levels (Fig. 6 A, B) in a similar magnitude as we observed it earlier in *Pclo*^*gt/gt*^ neurons (Fig. 3 F, G). Furthermore the presence of Rab5^S34N^ in *Pclo*^*wt/wt*^ neurons leads to lower levels of EEA1 at PI(3)P organelles as well as fewer FYVE-Rab5-EEA1 triple positive early endosomes (Fig. 6 B, D, F). However, Rab5^S34N^ expression does not alter the amount of Rab5 available at PI(3)P organelles and significantly increases the fraction of FYVE-Rab5 double positive organelles (Fig. 6 C, D, E). This stands in contrast to what we observed earlier in *Pclo*^*gt/gt*^ neurons. In neurons lacking Piccolo no functional complex of Rab5 on PI(3)P membranes is formed whereas in Rab5^S34N^ expressing neurons this complex is formed however the GTPase is locked in an inactive state causing it to stay on the membrane for longer time periods. Besides that, the observed similarities between *Pclo*^*gt/gt*^ and *Pclo*^*wt/wt*^ expressing Rab5^S34N^ neurons support the initial hypothesis that less active Rab5 could in part be responsible for the reduced early endosome numbers in *Pclo*^*gt/gt*^ neurons.

**Figure 6.**
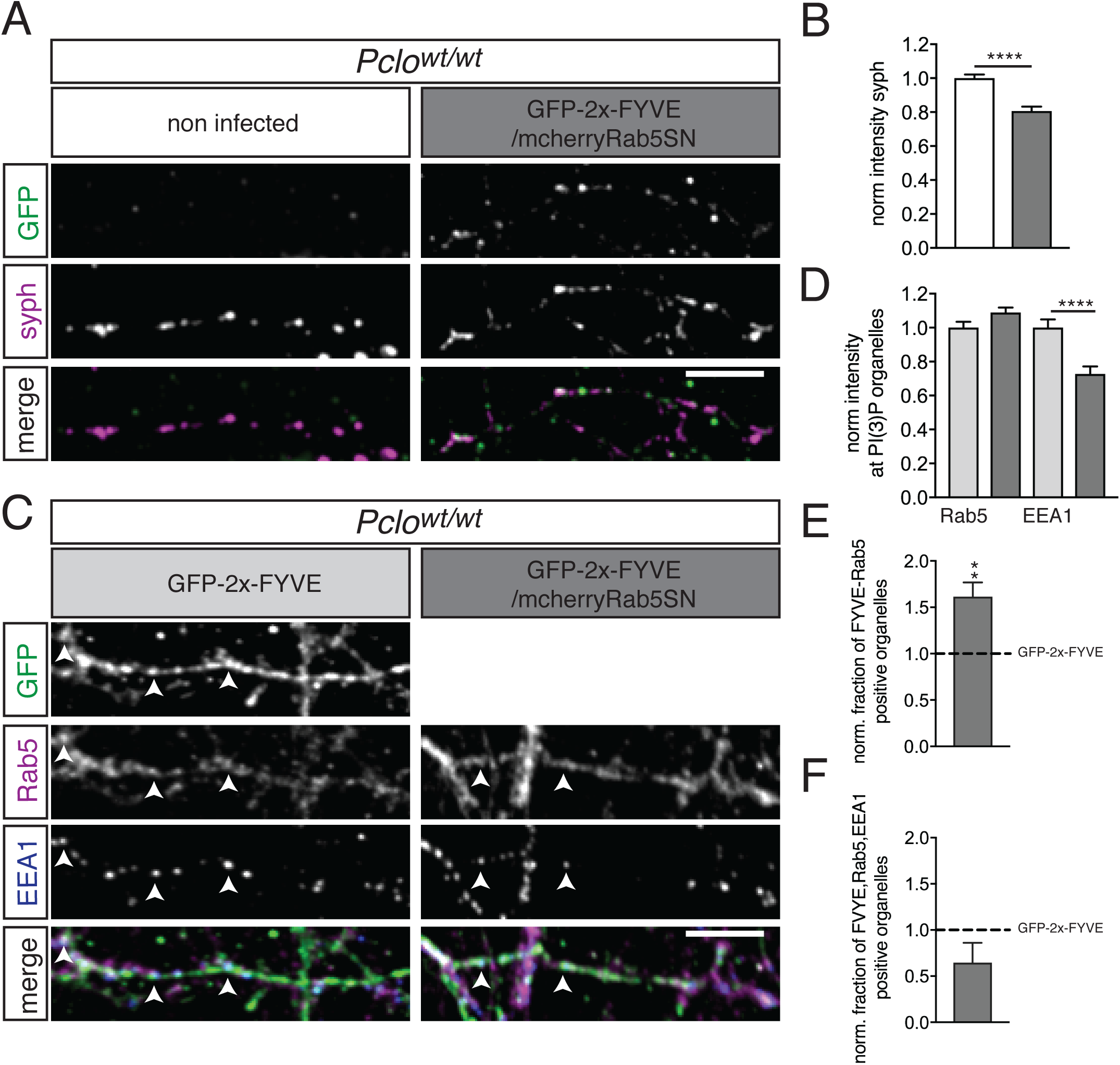
GDP-locked Rab5 (Rab5^S34N^) expression in *Pclo*^*wt/wt*^ neurons decreases Synaptophysin and early endosomes levels. (A, B) Rab5^S34N^ expression (GFP-2x-FYVE/mcherry-Rab5^S43N^) in *Pclo*^*wt/wt*^ neurons decreases Synaptophysin levels (*Pclo*^*wt/wt*^ = 1 ± 0.02, n = 214 synapses; *Pclo*^*wt/wt*^ + Rab5SN = 0.81 ± 0.03, n = 142 synapses; p < 0.0001; 2 independent experiments, students t-test). (C) Rab5 and EEA1 levels at PI(3)P organelles in neurons expressing Rab5^S34N^ (GFP-2x-FYVE/mcherry-Rab5^S43N^). (D) Quantification of C. Rab5^S34N^ expression increases Rab5 levels at PI(3)P organelles (*Pclo*^*wt/wt*^ = 1 ± 0.03, n = 930 puncta; *Pclo*^*wt/wt*^ + Rab5^S43N^ = 1.09 ± 0.03, n = 612 puncta; p = 0.06; 3 independent experiments, student`s t-test). EEA1 levels at PI(3)P organelles decrease upon Rab5^S34N^ expression (*Pclo*^*wt/wt*^ = 1 ± 0.05, n = 930 puncta; *Pclo*^*wt/wt*^ + Rab5^S34N^ = 0.73 ± 0.04, n = 613; p < 0.0001; 3 independent experiments, student t-test). (E) Quantification of C. More FYVE/Rab5 positive organelles are present in Rab5^S34N^ expressing neurons (*Pclo*^*wt/wt*^ = 1 ± 0; *Pclo*^*wt/wt*^ + Rab5^S34N^ = 1.614 ± 0.15; p = 0.007; 4 independent experiments). (F) Quantification of C. Fewer FYVE/Rab5/EEA1 triple positive organelles are present in Rab5^S43N^ expressing neurons (*Pclo*^*wt/wt*^ = 1 ± 0; *Pclo*^*wt/wt*^ + Rab5^S43N^ = 0.64 ± 0.22; p = 0.511; 4 independent experiments). Mean ± SEM; scale bar represents 10 μm, Students`s *t* - test.

### GTPase deficient Rab5 (Rab5^Q79L^) expression rescues EEA1 and Synaptophysin levels to *Pclo*^*wt/wt*^ amounts

As Rab5^S34N^ expression in *Pclo*^*wt/wt*^ neurons mimics *Pclo*^*gt/gt*^ phenotypes, we next tested whether the expression of GTPase deficient Rab5^Q79L^ can rescue the loss of Piccolo. This was accomplished by expressing Rab5^Q79L^ (Stenmark et al., 1994) tagged with mCherry (mCh-Rab5^Q79L^) together with GFP-2x-FYVE in *Pclo*^*wt/wt*^ and *Pclo*^*gt/gt*^ neurons. Analyzing Rab5 and EEA1 levels at PI(3)P organelles revealed a significant rescue of endogenous EEA1 levels in *Pclo*^*gt/gt*^ neurons (Fig. 7 A, B, D). Moreover, the fraction of early endosomes (FYVE/Rab5/EEA1) increased (Fig. 7, B and F), although the fraction of vesicles positive only for FYVE and Rab5 did not change (Fig. 7 E). Remarkably, the expression of Rab5^Q79L^ in *Pclo*^*wt/wt*^ neurons had the opposite effect, decreasing EEA1 levels (Fig. 7, A and D), though total Rab5 levels were not altered (Fig. 7, A and C). These data support the concept that the loss of Piccolo leads to altered Rab5 activity at synapses and along axons and therefore to a reduction in the maturation of early endosomes.

**Figure 7.**
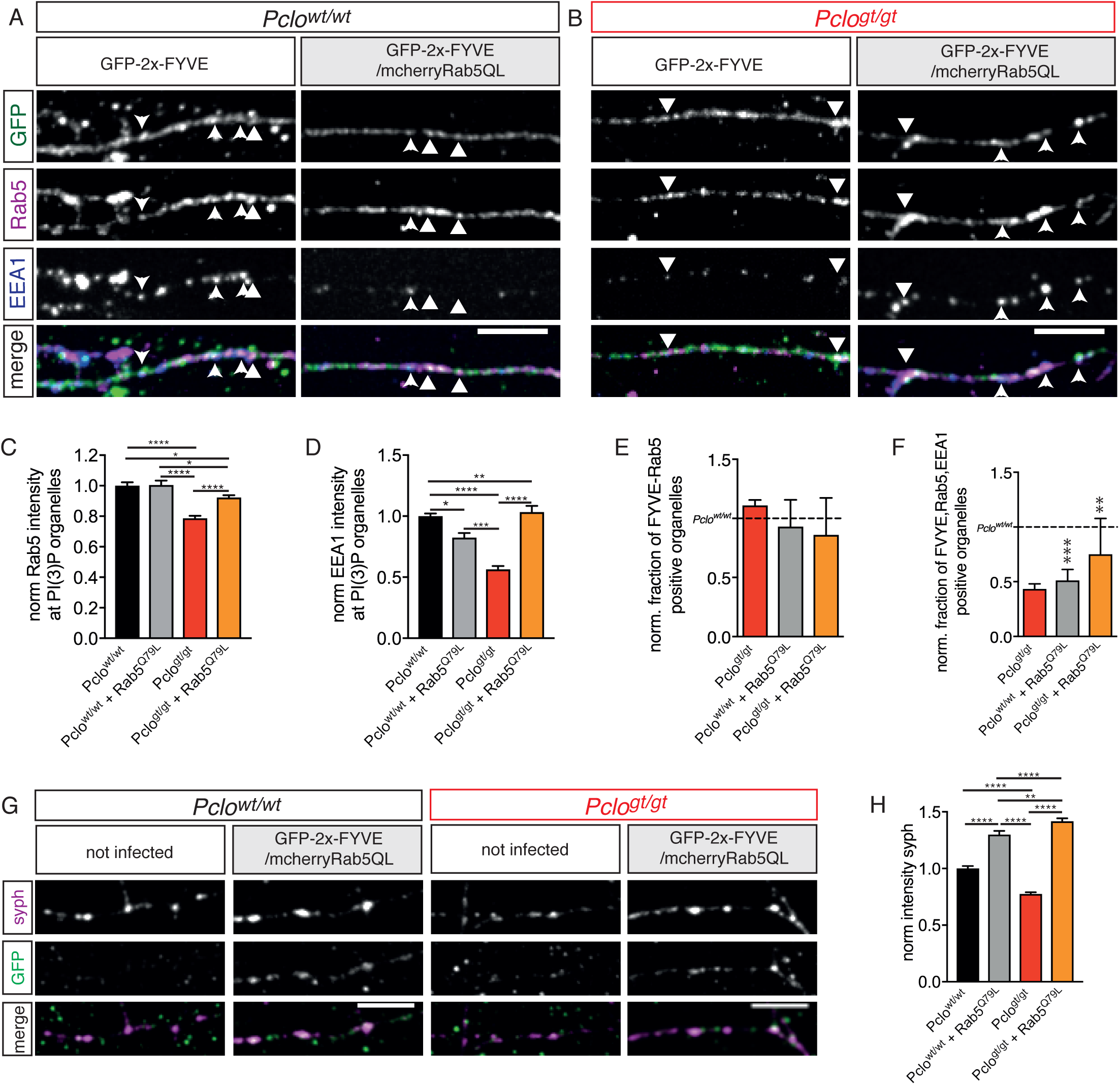
Expression of GTPase deficient Rab5 (Rab5^Q79L^) in *Pclo*^*gt/gt*^ neurons rescues EEA1 levels at PI(3)P membranes and Synaptophysin back to *Pclo*^*wt/wt*^ levels. (A) Rab5 and EEA1 levels at GFP-2x-FYVE organelles along axons in *Pclo*^*wt/wt*^ neurons expressing Rab5^Q79L^. (B) In *Pclo*^*gt/gt*^ neurons, Rab5^Q79L^ expression increases Rab5 and EEA1 level at GFP-2x-FYVE organelles towards *Pclo*^*wt/wt*^ levels. (C–F) Quantification of A and B. (C) Rab5^Q79L^ expression in *Pclo*^*wt/wt*^ neurons does not alter the total amount of Rab5 at GFP-2x-FYVE membranes (*Pclo*^*wt/wt*^ = 1 ± 0.01, n = 965 puncta; *Pclo*^*wt/wt*^ _(Rab5Q79L)_ = 1.00 ± 0.03, n = 980 puncta; p = 0.8983; 3 independent experiments). Rab5^Q79L^ expression in *Pclo*^*gt/gt*^ neurons does not increase total Rab5 levels at PI(3)P organelles (*Pclo*^*gt/gt*^ = 0.77 ± 0.01, n =840 puncta; *Pclo*^*gt/gt*^ _(Rab5Q79L)_ = 0.92 ± 0.01, n = 1185 puncta; p < 0.0001; 3 indepnedent experiments). (D) In *Pclo*^*wt/wt*^ neurons Rab5^Q79L^ expression causes EEA1 levels to drop (*Pclo*^*wt/wt*^ = 1 ± 0.03, n = 965 puncta; *Pclo*^*wt/wt*^ (Rab5Q79L) = 0.82 ± 0.04, n = 980 puncta; p = 0.0007; 3 independent experiments). Rab5^Q79L^ expression rescues EEA1 levels in *Pclo*^*gt/gt*^ neurons (*Pclo*^*gt/gt*^ = 0.56 ± 0.03, n = 840 puncta; *Pclo*^*gt/gt*^ _(Rab5Q79L)_ = 1.03 ± 0.05, n = 1183 puncta; p < 0.0001; 3 independent experiments). (E) The fraction of GFP-2x-FYVE/Rab5 double positive vesicles in *Pclo*^*wt/wt*^ and *Pclo*^*gt/gt*^ neurons is not changed upon Rab5^Q79L^ expression (*Pclo*^*gt/gt*^ = 1.11 ± 0.04, p = 0.0756; *Pclo*^*wt/wt*^ _(Rab5Q79L)_ = 0.93 ± 0.23, p = 0.7751, *Pclo*^*gt/gt*^ _(Rab5Q79L)_ = 0.86 ± 0.31, p = 0.6781; 3 independent experiments). (F) Rab5^Q79L^ expression decreases the fraction of GFP-2x-FYVE/Rab5/EEA1 triple positive vesicles in *Pclo*^*wt/wt*^ neurons but increases it in *Pclo*^*gt/gt*^ neurons (*Pclo*^*gt/gt*^ = 0.44 ± 0.04, p = 0.0002; *Pclo*^*wt/wt*^ _(Rab5Q79L)_ = 0.51 ± 0.10, p = 0.008; *Pclo*^*gt/gt*^ _(Rab5Q79L)_ = 0.75 ± 0.32, p = 0.4885; n = 3 independent experiments). (G) Rab5^Q79L^ expression in *Pclo*^*wt/wt*^ neurons increases Synaptophysin intensities. In *Pclo*^*gt/gt*^ neurons Rab5^Q79L^ expression rescues Synaptophysin levels higher than *Pclo*^*wt/wt*^ levels. (H) Quantification of G. Synaptophysin puncta intensity increases upon Rab5^Q79L^ expression in *Pclo*^*wt/wt*^ neurons. In *Pclo*^*gt/gt*^ neurons Rab5^Q79L^ expression rescues Synaptophysin levels higher than *Pclo*^*wt/wt*^ levels (*Pclo*^*wt/wt*^ = 1 ± 0.02, n = 443 synapses; *Pclo*^*gt/gt*^ = 0.78 ± 0.01, n = 700 synapses; *Pclo*^*wt/wt*^ _(Rab5Q79L)_ = 1.3 ± 0.03, n = 424 synapses; *Pclo*^*gt/gt*^ _(Rab5Q79L)_ = 1.46 ± 0.03, n = 682 synapses; 3 independent experiment). Scale bars represent 10 μm. Mean ± SEM. C, D and H ANOVA with Tukey multi comparison test, E, F Student`s *t* -test.

To assess whether this manipulation also rescued SV pool size in *Pclo*^*gt/gt*^ neurons, we immuno-stained mCh-Rab5^Q79L^/GFP-2x-FYVE expressing neurons with Synaptophysin antibodies. This analysis showed an increase in Synaptophysin levels at *Pclo*^*wt/wt*^ as well as *Pclo*^*gt/gt*^ boutons (Fig. 6, G and H), indicating that the loss of SVs in *Pclo*^*gt/gt*^ neurons is indeed caused by the impaired maturation of early endosomes and that Piccolo plays a role in this pathway.

### Silencing synaptic activity affects synaptic levels of endosome proteins

In non-neuronal cells, early endosomes are known to form during receptor-mediated endocytosis (Grant & Donaldson, 2009). At synapses, endocytosis is associated with synaptic activity as part of the fusion and recycling of SVs. Given that cultured hippocampal neurons are intrinsically active (Minerbi et al., 2009), we considered the possibility that this activity creates a pool of Rab5 positive early endocytic vesicles in *Pclo*^*gt/gt*^ boutons that subsequently only poorly matures into early endosomes. To test this concept, hippocampal cultures were treated with Tetrodotoxin (TTX) for 24 h and the synaptic levels of the endosome markers Rabex5, Rab5 and EEA1 were examined (Fig. 8). Interestingly in *Pclo*^*wt/wt*^ neurons, the block of synaptic activity causes the levels of all markers (Rab5, EEA1 and Rabex5) to drop (30, 30 and 10% respectively) (Fig. 8 A, C, D, F G, I). This is consistent with a role of synaptic activity in the formation of early endosomes. Similarly, silencing synapses in *Pclo*^*gt/gt*^ neurons decreased the levels of Rabex5 and Rab5 (by 10 and 30%), to a similar extent as what is seen in *Pclo*^*wt/wt*^ synapses (Fig. 8 B, C, E, F, H, I). However, the overall levels of Rabex5 in untreated synapses are decreased at *Pclo*^*gt/gt*^ synapses compared to *Pclo*^*wt/wt*^ neurons (Fig. 8 H, I). These data indicate that synaptic activity regulates the dynamic recruitment of Rabex5 and Rab5 at excitatory synapses. Surprisingly, while EEA1 levels were reduced by about 30% at *Pclo*^*wt/wt*^ synapses due to silencing synaptic activity (Fig. 8 C and F), in synapses lacking Piccolo, silencing increased the already reduced levels of EEA1 by ∼ 20% (Fig. 8 D and F). Together these data support the concept that proteins involved in the formation of early endosomes in presynaptic boutons are recruited into synapses in response to synaptic activity. They also indicate that the initial endocytic mechanism still operates in boutons lacking Piccolo, albeit EEA1 recruitment is impaired.

**Figure 8.**
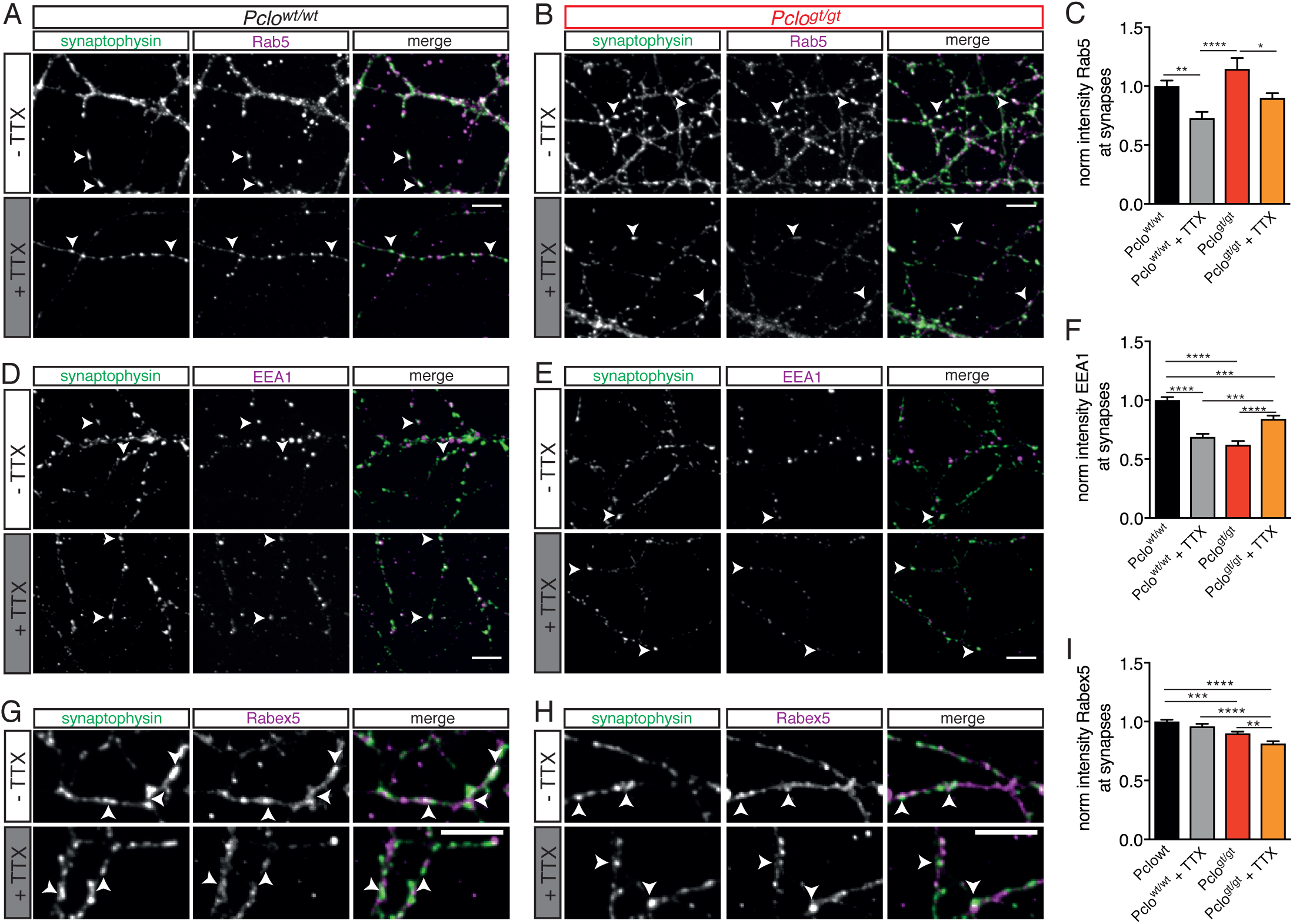
Silencing synaptic activity affects synaptic levels of endosome proteins in *Pclo*^*wt/wt*^ and *Pclo*^*gt/gt*^ neurons. (A) *Pclo*^*wt/wt*^ neurons stained for Synaptophysin and Rab5 after TTX treatment. (B).*Pclo*^*gt/gt*^ neurons stained for Synaptophysin and Rab5 after TTX treatment. (C) Quantification of A and B. Rab5 levels decrease in *Pclo*^*wt/wt*^ synapses upon TTX treatment (*Pclo*^*wt/wt*^ = 1 ± 0.05, n = 391 synapses; *Pclo*^*wt/wt*^ _(TTX)_ = 0.73 ± 0.05, n = 482 synapses; p < 0.01; 2 independent experiments). Synaptic Rab5 levels decrease in *Pclo*^*gt/gt*^ synapses (*Pclo*^*gt/gt*^ = 1.14 ± 0.09, n = 400 synapses; *Pclo*^*gt/gt*^ _(TTX)_ = 0.90 ± 0.04, n = 372 synapses; p < 0.05; 2 independent experiments). (D) *Pclo*^*wt/wt*^ neurons stained for Synaptophysin and EEA1 after TTX treatment. (E) *Pclo*^*gt/gt*^ neurons stained for Synaptophysin and EEA1 after TTX treatment. (F) Quantification of D and E. In *Pclo*^*wt/wt*^ synapses EEA1 level drop upon TTX treatment (*Pclo*^*wt/wt*^ = 1 ± 0.02, n = 1812 synapse; *Pclo*^*wt/wt*^ _(TTX)_ = 0.69 ± 0.02, n = 2034 synapses; p < 0.0001; 5 independent experiments). In contrast, EEA1 level increase in *Pclo*^*gt/gt*^ synapses due to TTX treatment (*Pclo*^*gt/gt*^ = 0.62 ± 0.03, n = 1119; *Pclo*^*gt/gt*^ _(TTX)_= 0.84 ± 0.03, n = 1581; p < 0.0001; 5 independent experiments). (G) *Pclo*^*wt/wt*^ neurons stained for Synaptophysin and Rabex5 after TTX treatment. (H) *Pclo*^*gt/gt*^ neurons stained for Synaptophysin and Rabex5 after TTX treatment. (I) Quantification of G and H. Rabex5 level at synapses decrease in *Pclo*^*wt/wt*^ and *Pclo*^*gt/gt*^ neurons upon TTX treatment (*Pclo*^*wt/wt*^ = 1 ± 0.01, n = 2150 puncta; *Pclo*^*wt/wt*^ _(TTX)_= 0.96 ± 0.02, n = 1906 puncta; *Pclo*^*gt/gt*^ = 0.90 ± 0.01, n = 2419, *Pclo*^*gt/gt*^ _(TTX)_ = 0.81 ± 0.02, n = 2277; p < 0.0001; 6 independent experiments). Scale bars represent 10 μm. Mean ± SEM, ANOVA with Tukey multi comparison test.

### Synaptic levels of Prenylated Rab Acceptor Protein 1 (Pra1) are reduced in *Pclo*^*gt/gt*^ neurons

We next asked the question how the active zone protein could be linked to the endosome pathway within presynaptic terminals. Interestingly Piccolo has been shown to interact with Pra1 (Fenster et al., 2000), which is a GDI replacement factor and is therefore part of the Rab-GTPase activating/deactivating cycle as it places the GTPase onto it`s target membrane (Pfeffer & Aivazian, 2004). Pra1, which is missing in KO neurons, could therefore be a possible explanation for the observed endosome phenotype in *Pclo*^*gt/gt*^ neurons. To test this hypothesis, we first analyzed the synaptic levels of Pra1 in *Pclo*^*wt/wt*^ and *Pclo*^*gt/gt*^ neurons. Here, we found that Pra1 localized to presynaptic terminals, co-localizing with Synaptophysin in *Pclo*^*wt/wt*^ neurons (Fig. 9 A). However, in *Pclo*^*gt/gt*^ neurons Pra1 levels were reduced at Synaptophysin positive synapses and accumulates in a spot like pattern outside synapses (Fig. 9 A, B, C). Together these data support suggest that the reduced formation of endosomal membranes in boutons lacking Piccolo is due to a reduced synaptic recruitment of it´s interaction partner Pra1.

**Figure 9.**
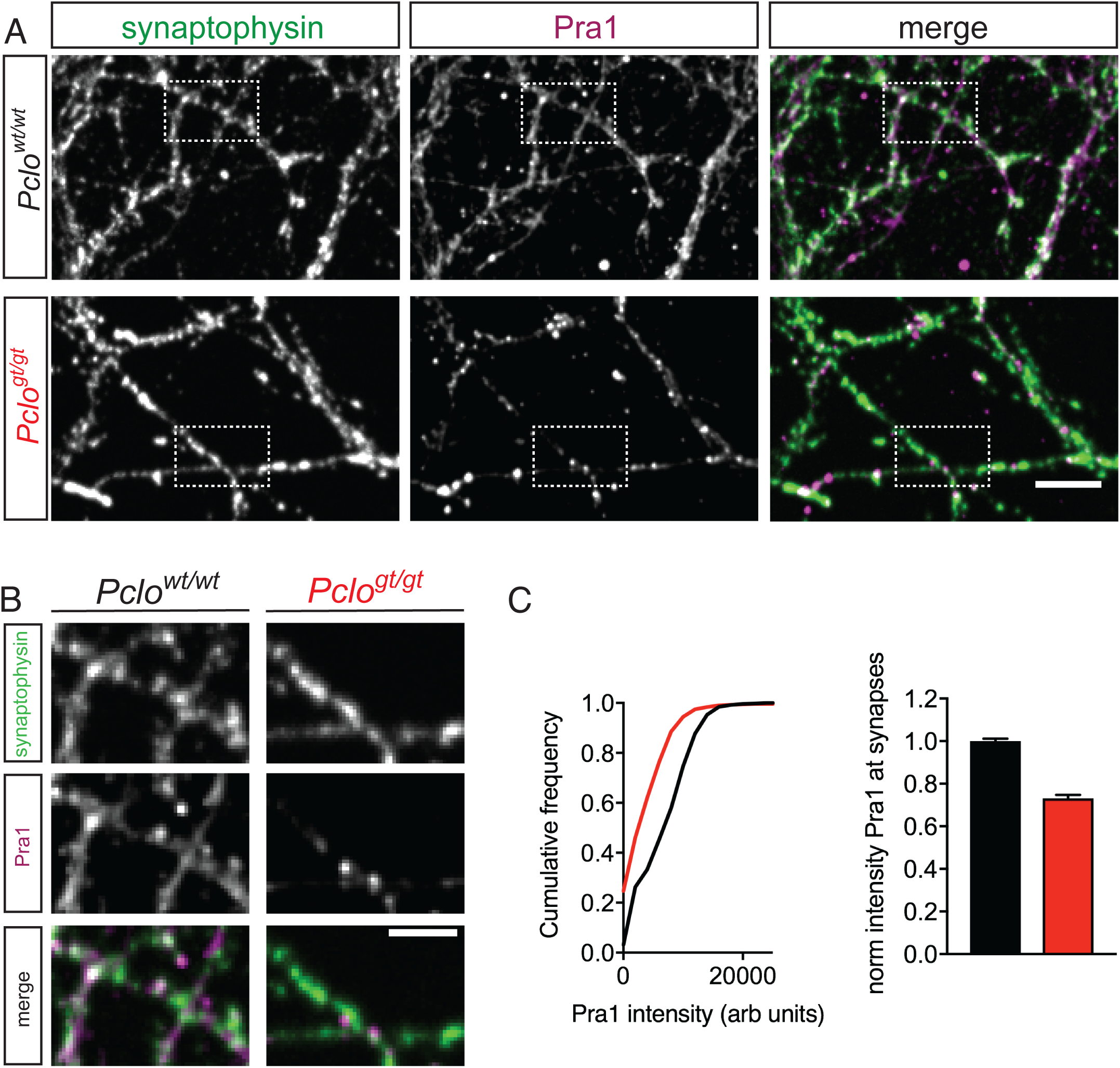
The loss of Piccolo leads to diminished levels of synaptic Pra1. (A) Pra1 is present at Synaptophysin positive synapses in *Pclo*^*wt/wt*^ synapses but is severely reduced in *Pclo*^*gt/gt*^ synapses. (B) Detail images of areas indicated in (A). (C) Cumulative frequency distribution of Pra1 intensity and normalized average Pra1 intensities in Synaptophysin puncta in *Pclo*^*wt/wt*^ and *Pclo*^*gt/gt*^ neurons. Loss of Piccolo decreases Pra1 intensity at synapses shifting its distribution to the left (left panel) (n = 1058 synapses for *Pclo*^*wt/wt*^, n = 1088 synapses for *Pclo*^*gt/g*t^, 3 independent experiments). Scale bar represents 10 μm and 5 μm in detailed image, Mean ± SEM, Student`s *t* –test.

### Expression of Pclo-Znf1 rescues synaptic Pra1 as well as EEA1 levels and synaptic vesicle pool size

As Piccolo interacts with Pra1 via its zinc fingers (Fenster et al., 2000), we hypothesized that this interaction is critical for its synaptic recruitment and subsequent role in the activation of Rab5 and the formation of early endosome. This suggests that the synaptic delivery of even one zinc finger could rescue the reduced synaptic levels of Pra1 and EEA1 in *Pclo*^*gt/gt*^ neurons. To test this hypothesis, we lentivirally expressed mCherry tagged Znf1 of Piccolo (Pclo-Znf1-mChr) in *Pclo*^*wt/wt*^ and *Pclo*^*gt/gt*^ primary hippocampal neurons. Neurons DIV 14 – 16 were then fixed and stained for Synaptophysin, Pra1 and EEA1. As observed earlier Synaptophysin levels, representing the total pool of SVs, is reduced in *Pclo*^*gt/gt*^ neurons compared to *Pclo*^*wt/wt*^ (Fig. 10 A and C). Lentiviral expression of Pclo-Znf1-mCh not only leads to its accumulation at Synaptophysin positive sites (Fig. 10 A) in both *Pclo*^*gt/gt*^ and *Pclo*^*wt/wt*^ neurons, but rescues Synaptophysin levels to greater than wildtype levels in *Pclo*^*gt/gt*^ and has only a modest effect in *Pclo*^*wt/wt*^ neurons (Fig. 10 A and D). Pclo-Znf1-mCh over-expression also restored Pra1 levels at *Pclo*^*gt/gt*^ synapses (Fig. 10 B and E). Intriguingly synaptic Pra1 levels in *Pclo*^*gt/gt*^ neurons expressing Pclo-Znf1-mCherry are even 50 % higher than in *Pclo*^*wt/wt*^ neurons. Of note *Pclo*^*wt/wt*^ synapses harboring Pclo-Znf1-mCh over-expression in *Pclo*^*wt/wt*^ synapses display about 30 % increased levels of synaptic Pra1 compared to synapses of un-infected *Pclo*^*wt/wt*^ neurons, indicating that the Piccolo Znf1 is both necessary and sufficient to recruits Pra1 into presynaptic boutons. As a more direct test of its role in the formation of presynaptic early endosome, we examined whether restoring Pra1 levels, through the over-expression of Pclo-Znf1, would rescue EEA1 levels in *Pclo*^*gt/gt*^ neurons back to *Pclo*^*wt/wt*^ levels. As shown in figure 10 C. Pclo-Znf1-mCh expression significantly increases synaptic EEA1 levels in *Pclo*^*gt/gt*^ synapses. The levels are increased compared to *Pclo*^*wt/wt*^ synapses but as high as in *Pclo*^*wt/wt*^ synapses expressing Pclo-Znf1-mCherry (Fig. 10 C and F) further indicating that the presence of the Pclo-Znf1 supports the accumulation of synaptic Pra1 and along with it EEA1.

**Figure 10.**
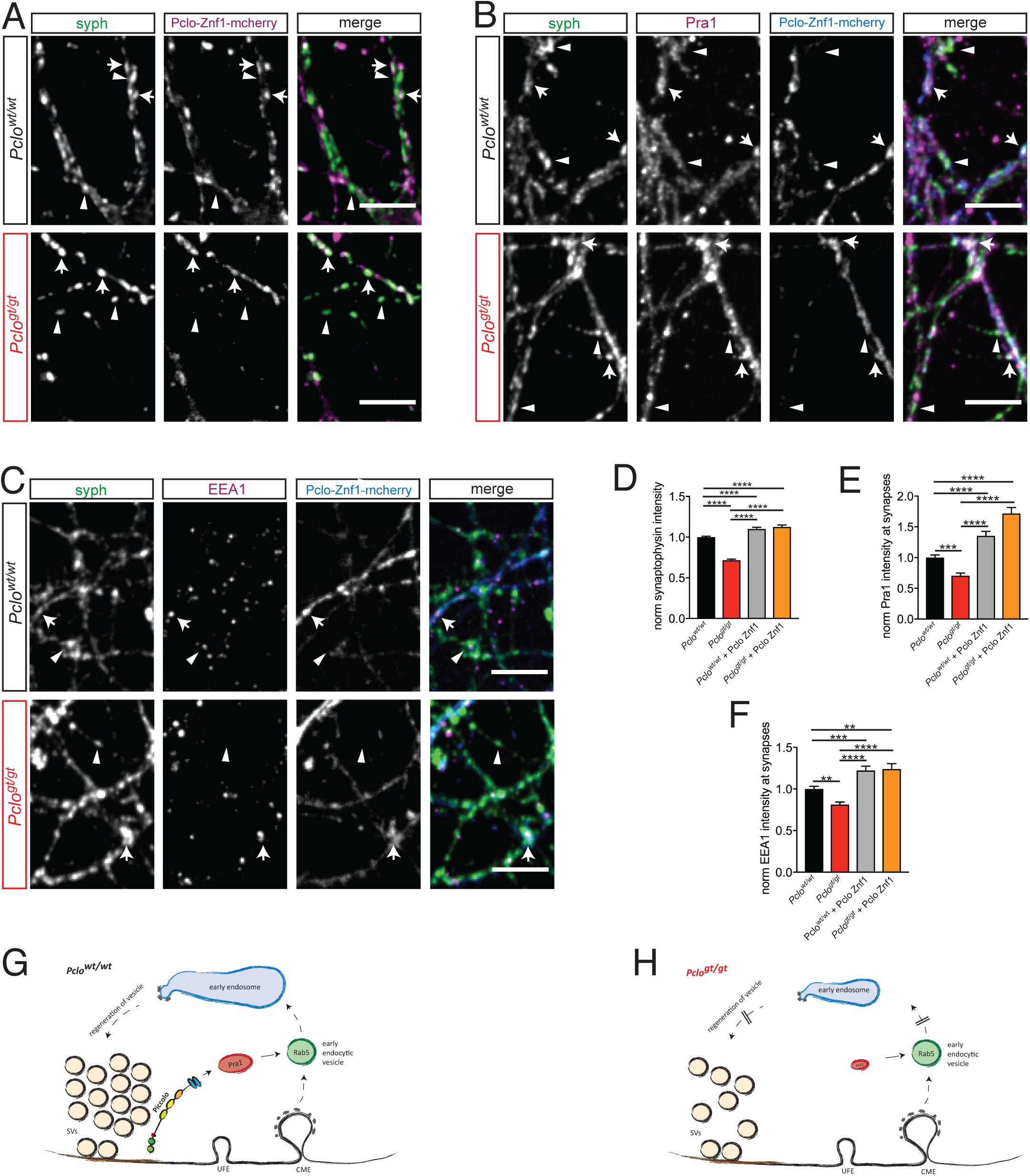
Expressing Pclo-Znf1-mCherry rescues Pra1, EEA1 and Synaptophysin levels in *Pclo*^*gt/gt*^ neurons. (A-C) *Pclo*^*wt/wt*^ and *Pclo*^*gt/gt*^ neurons expressing Pclo-Znf1-mCherry were fixed and stained for Synaptophysin (A), Pra1 (B) and EEA1 (C). (D) Quantification of A. ZnF1 slightly increases Synaptophysin levels in *Pclo*^*wt/wt*^ neurons (arrows vs arrowheads) (*Pclo*^*wt/wt*^ = 1 ± 0.01, n = 946 synapses; *Pclo*^*wt/wt*^ + Pclo-Znf1-mCherry = 1.1 ± 0.02, n = 391; p < 0.0001; 3 independent experiments). In Pclo-Znf1-mCherry expressing *Pclo*^*gt/gt*^ boutons (arrows), ZnF1 increases Synaptophysin levels compared to non-expressing boutons (arrowheads). (*Pclo*^*gt/gt*^ = 0.72 ± 0.01, n = 815; *Pclo*^*gt/gt*^ + Pclo-Znf1-mCherry = 1.13 ± 0.02, n = 387; p < 0.0001; 3 independent experiments). (E) Quantification of B. Pclo-Znf1-mCherry increases synaptic Pra1 levels in *Pclo*^*gt/gt*^ neurons (*Pclo*^*wt/wt*^ = 1 ± 0.04, n = 893 synapses; *Pclo*^*wt/wt*^ + Pclo-Znf1-mCherry = 1.35 ± 0.07, n = 494; p < 0.0001; 3 independent experiments). Pclo-Znf1-mCherry rescues and further increases synaptic Pra1 levels in *Pclo*^*gt/gt*^ neurons (*Pclo*^*gt/gt*^ = 0.71 ± 0.04, n = 724; *Pclo*^*gt/gt*^ + Pclo-Znf1-mCherry = 1.72 ± 0.10, n = 436; p < 0.0001; 3 independent experiments). (F) Quantification of C. Pclo-Znf1-mCherry increases synaptic EEA1 levels in *Pclo*^*wt/wt*^ neurons (*Pclo*^*wt/wt*^ = 1 ± 0.03, n = 1053 synapses; *Pclo*^*wt/wt*^ + Pclo-Znf1-mCherry = 1.22 ± 0.05, n = 709; p < 0.0001; 3 independent experiments). Pclo-Znf1-mCherry rescues and further increases synaptic EEA1 levels in *Pclo*^*gt/gt*^ neurons (*Pclo*^*gt/gt*^ = 0.81 ± 0.03, n = 955; *Pclo*^*gt/gt*^ + Pclo-Znf1-mCherry = 1.24 ± 0.06, n = 464; p < 0.0001; 3 independent experiments). (G and H) Model describing the contribution of Piccolo in synaptic vesicle recycling. Piccolo regulates Rab5 function and subsequently early endosome formation through it`s interaction with Pra1. In the absence of Piccolo less Pra1 is localized within the presynaptic terminal. Less synaptic Pra1 negatively impacts Rab5 function and early endosome formation with the consequence that SVs are lost over time. Scale bars represent 10 μm. Mean ± SEM, ANOVA with Tukey multi comparison test.

In sum, these data support our hypothesis that Piccolo via its zinc fingers and its binding partner Pra1 plays critical roles during the recycling and maintenance of SV pools, acting through their activation of Rab5 and EEA1 and the formation of early endosomes (Fig 10 G and H).

## Discussion

Our study has shown that Piccolo plays a critical role in the activity dependent recycling of SVs. Specifically, we find that Piccolo loss of function reduces SV numbers as well as the recycling of SVs, without affecting the activity dependent docking and fusion of SVs. Mechanistically, we find that boutons lacking Piccolo accumulate endocytic vesicles that fail to mature into early endosomes due to an impaired Rab5 activation and recruitment of EEA1 towards PI(3)P positive membranes. This is in consequence of reduced presynaptic levels of Pra1. The phenotype can be restored by the over-expression of Rab5^Q79L^ as well as the ZnF domain of Piccolo, which interacts with and restores Pra1 levels in Piccolo deficient synapses.

### Ultrastructural changes in boutons lacking Piccolo

Piccolo and its isoforms are encoded by a large 380 kb gene (*Pclo)* (Fenster & Garner, 2002). Using transposon mediated mutagenesis (Izsvak et al., 2010), a rat line was created that disrupts the downstream expression of exons 4-25 (Medrano et al., 2018). Western blotting of brain lysate reveals a near complete loss of all dominant high and low molecular weight isoforms of Piccolo (Fig. 1 C). This is associated with an almost complete synaptic loss of Piccolo immuno-reactivity (Fig. 1). However the loss of Piccolo does not affect the formation of synapses (Fig. 1 and 2), which is consistent with previous studies (Leal-Ortiz et al., 2008; Mukherjee et al., 2010). Intriguingly, our quantitative analysis revealed a large decrease in the number of SVs/bouton (Fig. 2). This was associated with a concomitant increase in the number of small endocytic-like vesicles (∼80 nm), suggesting that Piccolo is important for the recycling of SVs. It is worth noting, that the deletion of exon 14 in the mouse *Pclo*^*KO/KO*^ gene, which appears to only remove the longest Piccolo isoforms (Mukherjee et al., 2010; Waites et al., 2011), does not appear to affect the number of SVs/bouton or lead to an increase in the steady-state levels of endocytic structures (Mukherjee et al., 2010). The continued expression of lower molecular weight variant retaining ZnF1 and/or ZnF2 and thus the synaptic recruitment of Pra1 could explain this difference (see below).

### Synaptic transmission in boutons lacking Piccolo

A fundamental question is how the loss of Piccolo and the reduced number of SVs adversely impacts synapse function? Our electrophysiological data indicate that the evoked release of neurotransmitter is not affected. This is somewhat surprising as Piccolo has been shown to physically interact with AZ proteins including RIM, RimBP, VGCC and Munc13 (Gundelfinger et al., 2015), which are known to be involved in SV priming and calcium mediated SV fusion (Breustedt et al., 2010; Davydova et al., 2014; Girach, Craig, Rocca, & Henley, 2013; Schoch et al., 2002). However, a functions in synaptic release could easily be masked by complementary roles played by other AZ proteins such as Bassoon (Gundelfinger et al., 2015).

Although RRP, Pvr and PPR were not altered (Fig. 3), we did observe an enhanced rundown of EPSC amplitudes during a 10 Hz stimulus train in boutons lacking Piccolo (Fig. 3). Conceptually, this could be related to a faster depletion of a smaller reserve pool of SVs or some defect in the retrieval of SVs from the plasma membrane. Reduced levels of FM4-64 dye uptake following stimulation in *Pclo*^*gt/gt*^ boutons support the idea that the TRP of SVs is smaller (Fig. 3). Furthermore, the larger number of endocytic-like vesicles (Fig. 2) suggests that the loss of Piccolo impairs efficient recycling of SVs, reducing the pool of SVs. Consistent with this concept, the destaining rate of FM1-43 during a 10 Hz 900 AP stimulation in boutons lacking Piccolo was dramatically slowed indicating that SVs are not efficiently regenerated within this time frame (Fig. 3)

### Early endosome trafficking is attenuated in boutons lacking Piccolo

An important question is how Piccolo loss of function impairs SV regeneration/recycling. Interestingly, the loss/inactivation of endocytic proteins, like Endophilin, Synaptojanin, Dynamin and the small GTPase Rab5, cause similar phenotypes to those observed in *Pclo*^*gt/gt*^ neurons (Hayashi et al., 2008; Milosevic et al., 2011; Schuske et al., 2003; Wucherpfennig et al., 2003). Of these, Endophilin and Dynamin are involved in the direct uptake of vesicles from the plasma membrane through endocytosis. However, our data with GFP-2x-FYVE argues against a block in early events of endocytosis in *Pclo*^*gt/gt*^ synapses, as a) eGFP-2x-FYVE levels are not significantly altered in Piccolo KO neurons (Fig. 4) and b) PI(3)P is added to already pinched off vesicles after the dephosphorylation of Phosphatidylinositol-4,5-bisphosphate (PI(4,5)P2) via Synaptojanin (Harris, Hartwieg, Horvitz, & Jorgensen, 2000; Mani et al., 2007; Verstreken et al., 2003). Importantly, GFP-2x-FYVE organelles are smaller in *Pclo*^*gt/gt*^ boutons (Fig. S3). This indicates that while the initial uptake of vesicle proteins is not affected, the subsequent formation of the early endosome compartment is disturbed. Of note, our EM analysis revealed the accumulation of round vesicles with a diameter > 60 nm, which could represent early endocytic vesicles (Fig. 2).

In line with this concept, we observed that loss of Piccolo differentially affected the appearance of early and late endosomes. For example, Rab5 vesicles accumulate along *Pclo*^*gt/gt*^ axons, whereas EEA1 and Rab7 vesicles decrease (Fig. 4) indicating that the loss of Piccolo leads to an early pre EEA1 block in the maturation of the endosome compartment. Importantly, the lower levels of Rab7 indicate that the trafficking of proteins through late endosomal structures could be affected, which subsequently affects the efficient degradation of defective molecules, a topic worth exploring in future studies. PI(3)P as well as Rab5-GTP are necessary to promote EEA1 mediated fusion of early endocytic vesicles (Spang, 2009). PI(3)P levels are not altered in *Pclo*^*gt/gt*^ neurons (Fig. 4), suggesting a possible defect in the activation/stabilization of Rab5-GTP. Normally, Rab5 is activated, amongst others, via its GEF Rabex5 (Horiuchi et al., 1997). Interestingly, besides reduced levels of Rabex5 at PI(3)P sites, fewer organelles harboring a complex of Rabex5 and Rab5 were observed in *Pclo*^*gt/gt*^ neurons (Fig. 5), a complex necessity for the recruitment of EEA1 and the formation of early endosomes (Murray et al., 2016; Simonsen et al., 1998; Stenmark, Vitale, Ullrich, & Zerial, 1995). This observation indicates that less active Rab5 is present in the boutons of *Pclo*^*gt/gt*^ neurons, a concept further supported by the fact that the expression of a GDP-locked Rab5 (Rab5^S34N^) in *Pclo*^*wt/wt*^ neurons also causes a reduction of EEA1 at PI(3)P organelles and therefore fewer early endosomes (Fig. 6). However Rab5^S34N^ expression does lead to more FYVE-Rab5 double positive organelles indicating that the inactive GTPase still becomes associated with PI(3)P organelles. This is different in *Pclo*^*gt/gt*^ neurons, where less Rab5 is present at PI(3)P organelles indicating that the loss of Piccolo may cause a destabilization of Rab5 on endosome membranes (Fig 5, 6). Consistently, a constitutive active Rab5 (Rab5^Q79L^) rescued EEA1 level at PI(3)P organelles as well as early endosome numbers in *Pclo*^*gt/gt*^ neurons (Fig. 7). It also rescued Synaptophysin back to WT levels, indicating that Rab5 dependent early endosome formation contributes to SV pool size at excitatory vertebrate synapses (Fig. 7). This concept is consistent with studies in *Drosophila*, where Rab5 dominant negative constructs also block the maturation of early endosomes at the neuromuscular junction (NMJ), while the over-expression of WT Rab5 increases SV endocytosis and quantal content (Wucherpfennig et al., 2003). Furthermore, these data indicate that SVs are at least partially reformed through an early endosomes in Piccolo lacking boutons which is also reflected by the slower FM destain kinetics in *Pclo*^*gt/gt*^ neurons (Fig 3).

### The formation of early endosomes depends on synaptic activity

Although the molecular mechanisms regulating SV endocytosis have been studied in great detail (Saheki & De Camilli, 2012), less is known about the subsequent endocytic steps and their relationship to synaptic transmission and SV recycling. Important open questions include whether the early endosome compartment is part of the SV cycle and thus activity dependent? At present, the former issue is still hotly debated. For example, Hoopmann and colleagues could show that SV proteins travel through an endosome compartment, while they are recycled (Hoopmann et al., 2010). Furthermore, it has been shown at the *Drosophila* NMJ that the synaptic FYVE-positive compartment disappears upon block of synaptic activity (Wucherpfennig et al., 2003). Consistently, we could show in rat hippocampal neurons that synaptic levels of endosome proteins (Rab5, EEA1 & Rabex5) significantly decrease upon TTX treatment (Fig. 8), indicating that early endosome formation in vertebrate synapses is also activity dependent. A decrease upon TTX treatment is also seen in *Pclo*^*gt/gt*^ synapses, although only for Rabex5 and Rab5, but not for EEA1, which is already low (Fig. 8). Taken together these data show that presynaptic early endosome formation depends on synaptic activity in *Pclo*^*wt/wt*^ neurons and in part also in *Pclo*^*gt/gt*^ neurons.

### Links between Piccolo and the endosome pathway

Alongside activity, our data show that Rab5 function and subsequently the recruitment of EEA1 towards PI(3)P membranes, also depends significantly on the presence of the active zone protein Piccolo (Fig. 6). As discussed above, Rab5 function is amongst others regulated by the GEF Rabex5, which forms a complex together with Rabaptin-5 (Horiuchi et al., 1997; Lippe, Miaczynska, Rybin, Runge, & Zerial, 2001; Mattera, Tsai, Weissman, & Bonifacino, 2006; Stenmark et al., 1995). Interestingly, Rabex5 and Rabaptin-5 loss of function studies in *C. elegans*, also affecting Rab5 function, show similar phenotypes to what we observe, e.g. fewer endosomes and less Synaptophysin immuno-reactivity (Sann, Crane, Lu, & Jin, 2012). However our data indicate that it is the interaction between Piccolo and the GDI replacement factor (GDF) Pra1 that promotes Rab5 function at vertebrate synapses. Previous studies have shown that Piccolo can directly interact with Pra1 (Fenster et al., 2000) and that Pra1 promotes the recruitment of Rab5-GDP to endosomal membranes (Abdul-Ghani, Gougeon, Prosser, Da-Silva, & Ngsee, 2001; Fenster et al., 2000; Hutt, Da-Silva, Chang, Prosser, & Ngsee, 2000; McLauchlan et al., 1998). Pra1 is a suitable target to regulate Rab5 function as it, among others, interacts with it and other endosomal GTPases (Pfeffer & Aivazian, 2004). As Pra1 nicely co-localizes with Synaptophysin, a marker for presynaptic terminals (Fig. 9 A), it is likely to target cytosolic GDI bound Rab5 towards endosome membranes in the presynapse. Membrane targeting is a requirement for the successive activation of GTPases by their designated GEF (Cherfils & Zeghouf, 2013). In line with this concept, synaptic levels of Pra1 are reduced in *Pclo*^*gt/gt*^ synapses, which could contribute to lower Rab5 levels on endosome membranes and consequently to reduced EEA1 levels and the formation of fewer early endosomes (Fig, 6). Importantly, the expression just the Piccolo Znf1 domain, known to bind Pra1 (Fenster et al., 2000), in *Pclo*^*gt/gt*^ neurons was sufficient to restore/increase presynaptic levels of both Pra1 and EEA1 to greater than WT levels (Fig. 10). This data support our hypothesis that Piccolo promotes the localization & stabilization of Pra1 within presynaptic boutons and thus the Rab5/EEA1 dependent formation of early endosomes (Fig. 10 G and H).

Finally, it is worth considering the disease relevance of the observed changes at *Pclo*^*gt/gt*^ synapses. As mentioned above, Piccolo has been implicated in psychiatric, developmental and neurodegenerative disorders. Although Piccolo is linked to all of these diseases, the molecular mechanism and its contribution to these disorders is unclear. The current study provides insights into one possible underlying mechanism. Specifically, the observed defects in early endosome formation are anticipated to affect the normal trafficking of SV proteins and membranes, a situation that could adversely affect the functionality of synapses throughout the brain, by impairing synaptic transmission, the maintenance of SV pools and the integrity of synapses. Anatomically, these alterations could destabilize synapses leading, for example, to neuronal atrophy during aging, a concept consistent with recent studies showing that Piccolo along with Bassoon regulates synapse integrity through the endo-lysosome and autophagy systems (Okerlund et al., 2017; Waites et al., 2013). Future studies will help clarify these relationships.

## Materials and Methods

### Hippocampal cell culture preparation

Microisland cultures were prepared from hippocampal neurons and maintained as previously described (Arancillo et al., 2013). All procedures for experiments involving animals, were approved by the animal welfare committee of Charité Medical University and the Berlin state government. Hippocampi were harvested from *Pclo*^*wt/wt*^ and *Pclo*^*gt/gt*^ (Wistar) P0-2 rats of either sex. Neurons were plated at 3000 cells/35 mm well on mouse astrocyte microislands to generate autaptic neurons for electrophysiology experiments.

For live cell imaging and immunocytochemistry, hippocampal neuron cultures were prepared using the Banker culture protocol (Banker & Goslin, 1988; Meberg & Miller, 2003; Tanaka, 2002). Astrocytes derived from mouse P0-2 cortices were plated into culture dishes 5-7 d before adding neurons. Nitric acid treated coverslips with paraffin dots were placed in separat culture dishes and covered with complete Neurobasal-A containing B-27 (Invitrogen, Thermo Fisher scientific, Waltham, USA), 50 U/ml Penicillin and 50 μg/ml streptomycin (Invitrogen, Thermo Fisher scientific, Waltham, USA). *Pclo*^*wt/wt*^ and *Pclo*^*gt/gt*^ hippocampi were harvested from P0-2 brains in ice cold HBSS (Gibco, Thermo Fisher scientific, Waltham, USA) and incubated consecutive in 20 U/ml papain (Worthington, Lakewood, USA) for 45-60 min and 5 min in DMEM consisting of albumin (Sigma-Aldrich, St. Louis, USA), trypsin inhibitor (Sigma-Aldrich, St.Louis, USA) and 5 % FCS (Invitrogen, Thermo Fisher scientific, Waltham, USA) at 37°C. Subsequent tissue was transferred to complete Neurobasal-A, and triturated. Isolated cells were plated on coverslips with paraffin dots at a density of 100,000 cells/35 mm well and 50,000 cells/20 mm well. After 1,5 h coverslips were flipped upside down and transferred to culture plates containing astrocytes in complete Neurobasal-A. When necessary, neurons were transduced with lentiviral constructs 72-94 h after plating. Cultures were incubated at 37 °C, 5% CO_2_ for 10-14 days before starting experiments.

### Genotyping Pclo rats

Ear pieces taken from rats were digested over night at 50°C in SNET-buffer (400mM NaCl, 1% SDS, 200 mM Tris (pH 8.0), 5 mM EDTA) containing 10 mg/ml proteinase K. Subsequently samples were incubated for 10 min at 99 °C and centrifuged for 2 min at 13.000 rpm. DNA was precipitated from the supernatant with 100 % isopropanol, centrifuged for 15 min at 13.000 rpm, washed with 70 % ethanol and centrifuged again for 10 min at 13.000 rpm. The obtained pellet was re-suspended in H_2_O. PCR reaction with a specific primer combination was performed to determine genotypes. The following primers were used: F2: 5`gcaggaacacaaaccaacaa3`; R1: 5` tgacctttagccggaactgt3`; SBF2: 5`tcatcaaggaaaccctggac3`. The PCR reaction protocol was the following: 2 min 94 °C; 3 × (30 sec 94 ° C, 60 ° C 30 sec, 72 ° C 30 sec); 35 × (94 ° C 30 sec, 55 ° C 30 sec, 72 ° C 30 sec); 72 ° C 10 min.

### Plasmid constructions

Rab5 and Rab7 were obtained from Richard Reimer (Stanford University) and subcloned inframe with the C-terminus of GFP as BsrG1-EcoR1 fragments in the lentiviral vector FUGWm (Leal-Ortiz et al., 2008). Third generation lentiviral vectors were generated using plasmids previously described (Campeau et al., 2009). GFP-2x-FYVE (Gillooly et al., 2000) was cloned into an ENTRY vector and transferred into pLenti-CMV-Neo-Dest (gift from Eric Campeau, Addgene plasmid # 17392) by a gateway LR recombination. For co-expression of GFP-2x-FYVE with mCherry-Rab5 (Q79L), we generated a bi-cistronic expression cassette linking GFP-2x-FYVE and mCherry-Rab5 (Q79L) by an IRES sequence. The resulting ENTRY vectors were transferred by Gateway LR recombination into a gateway-enabled destination vector derived from pCDH-EF1a-MCS-IRES-PURO (SystemBiosciences), which lacked the IRES sequence and the puromycin resistance cassette. For co-expression of GFP-2x-FYVE with mCherry-Rab5 (S34N), we generated a bi-cistronic expression cassette linking GFP-2x-FYVE and mCherry-Rab5 (S34N) by an IRES sequence. The resulting ENTRY vectors were transferred by Gateway LR recombination into a gateway-enabled destination vector derived from pCDH-EF1a-MCS-IRES-PURO (SystemBiosciences), which lacked the IRES sequence and the puromycin resistance cassette.

### Lentivirus production

Lentivirus was produced as previously described (Haferlach & Schoch, 2002). In brief, an 80% confluent 75 cm^2^ flask of HEK293T cells was transfected with 10 µg shuttle vector and mixed helper plasmids (pCMVd8.9 7.5 µg and pVSV-G 5 µg) using XtremeGene 9 DNA transfection reagent (Roche Diagnostics, Mannheim, Germany). After 48 h, cell culture supernatant was harvested and cell debris was removed by filtration. Aliquots of the filtrate were flash frozen in liquid nitrogen and stored at -80°C until use. Viral titer was estimated by counting cells in mass culture WT hippocampal neurons expressing GFP or mCherry as fluorescent reporter. Primary hippocampal cultures were infected with 80 µl of the viral solution (0.5-1 × 10^6^ IU/ml) 72-96 h post-plating.

### Cell Transfection with Lipofectamine

Sixty % confluent HEK293 cells were transfected with lipofectamine 2000 as described by the manufacturer (Invitrogen, Thermo Fisher scientific, Waltham, USA). In brief, 500 µl Optimem (Invitrogen, Thermo Fisher scientific, Waltham, USA) and 1 μg plasmid DNA were mixed and incubated for at least 5 min before adding 500 μl Optimem containing 15 µl lipofectamine 2000 (Invitrogen, Thermo Fisher scientific, Waltham, USA). After 20 min the mixture was added to cells.

### Western blot analysis

Brains from P0–2 animals as well as hippocampal neurons 14 days in vitro (DIV) were lysed in Lysis Buffer (50 mM Tris-HCl, 150 mM NaCl, 5 mM EDTA, 1 % TritonX-100, 0.5 % Deoxycholate, protease inhibitor pH 7.5) and incubated on ice for 5 min. Samples were centrifuged at 4°C with 13.000 rpm for 10 min, supernatant was transferred into a fresh tube and the protein concentration was determined using a BCA protein assay kit (Thermo Fisher scientific, Waltham, Massachusetts, USA). The same protein amounts were separated by SDS-Page and transferred onto nitrocellulose membranes (running buffer: 25 mM Tris, 190 mM glycin, 0.1 % SDS, pH 8.3; transfer buffer: 25 mM Tris, 192 mM Glycine, 1% SDS, 10% Methanol for small proteins, 7% Methanol for larger proteins pH 8.3). Afterwards membranes were blocked with 5 % milk in TBST (20 mM Tris pH 7.5, 150 mM NaCl, 0.1 % Tween 20) and incubated with primary antibodies in 3% milk in TBST o.n at 4°C. The following antibodies were used: Piccolo (1:1000; rabbit; synaptic systems, Göttingen, Germany; Cat# 142002, RRID:AB_887759), Piccolo (1:1000; rabbit; abcam, Cambridge, UK; Cat# ab20664, RRID:AB_777267), Piccolo (1:1000, guinea pig, synaptic systems, Göttingen, Germany; Cat# 142104, RRID:AB_2619831), Synaptophysin (1:1000; mouse; synaptic systems, Göttingen, Germany; Cat# 101011, RRID:AB_887824), VGlut1 (1:1000; guinea pig; synaptic systems, Göttingen, Germany; Cat# 135304, RRID:AB_887878), Rab5 (1:1000; mouse; synaptic systems, Göttingen, Germany; Cat# 108011, RRID:AB_887773), Rab7 (1:1000; rabbit; abcam, Cambridge, UK; Cat# ab137029, RRID:AB_2629474), EEA1 (1:1000; rabbit; cell signaling, Danvers, USA; Cat# 3288S, RRID:AB_2096811), Actin (1:1000; rabbit; Sigma-Aldrich, St. Louis, USA; Cat# A2066, RRID:AB_476693), Bassoon (1:1000; rabbit; synaptic systems, Göttingen, Germany; Cat# 141002, RRID:AB_887698), Homer (1:1000; guinea pig; synaptic systems, Göttingen, Germany; Cat# 160004, RRID:AB_10549720), PSD95 (1:5;mouse; NeuroMab, Davis, USA; Cat# 73-028, RRID:AB_10698024), Dynamin (1:1000; rabbit; abcam, Cambridge, UK; Cat# ab52611, RRID:AB_869531). The following day membranes were washed 3 times with TBST and incubated with HRP labeled secondary antibodies for 1 h at RT (Thermo Fisher scientific, Waltham, USA, dilution 1:1000). Subsequently membranes were washed 3 times with TBST and secondary antibody binding was detected with ECL Western Blotting Detection Reagents (Thermo Fisher scientific, Waltham,Massachusetts, USA) and a Fusion FX7 image and analytics system (Vilber Lourmat).

### Electrophysiology

Whole-cell voltage-clamp experiments were performed using one channel of a MultiClamp 700B amplifier (Molecular Devices, Sunnyvale, USA) under control of a Digidata 1440A Digitizer (Molecular Devices, Sunnyvale, USA) and pCLAMP Software (Molecular Devices, Sunnyvale, USA). Neurons were recorded at DIV 12-18. Neurons were clamped at -70 mV, EPSCs were evoked by 2 ms depolarization to 0 mV, resulting in an unclamped AP. Data were sampled at 10 kHz and Bessel filtered at 3 kHz. Series resistance was typically under 10 MΩ. Series resistance was compensated by at least 70%. Extracellular solution for all experiments, unless otherwise indicated, contained (in mM): 140 NaCl, 2.4 KCl, 10 HEPES, 10 glucose, 4 MgCl_2_, 2 CaCl_2_ (pH 7.4). Intracellular solution contained the following (in mM): 126 KCl, 17.8 HEPES, 1 EGTA, 0.6 MgCl_2_, 4 MgATP, 0.3 Na_2_GTP, 12 creatine phosphate, and phosphocreatine kinase (50 U/mL) (300 mOsm, pH7.4). All reagents were purchased from Carl Roth GMBH (Essen, Germany) with the exception of Na_2_ATP, sodium Na_2_GTP, and creatine-phosphokinase (Sigma-Aldrich, St. Louis, MO), and Phosphocreatine (EMD Millipore Chemicals, Billerica, USA).

Electrophysiological recordings were analyzed using Axograph X (Axograph, Berkley, USA), Excel (Microsoft, Redmond, USA), and Prism software (GraphPad, La Jolla, USA). EPSC amplitude was determined as the average of 5 EPSCs at 0.1 Hz. RRP size was calculated by measuring the charge transfer of the transient synaptic current induced by a 5 s application of hypertonic solution (500 mM sucrose in extracellular solution). Pvr was determined as a ratio of the charge from evoked EPSC and the RRP size of the same neuron. Short-term plasticity was analyzed evoking 50 synaptic responses at 10 Hz. PPR was measured dividing the second EPSC amplitude with the first EPSC amplitude elicited with an inter-pulse-interval of 25 ms.

### Electron microscopy

For electron microscopy, neurons were grown on 6-mm sapphire disks and processed as earlier described (Watanabe et al., 2013). In brief, cells were sandwiched between spacers and fixed through high-pressure freezing. Following freeze substitution with 1 % osmium and 1 % uranylacetate in acetone, sapphire discs were washed 4 times with acetone and postfixed with 0.1 % uranylacetate for 1 hour. Following 4 washing steps with acetone, cells were infiltrated with plastic (Epon, Sigma-Aldrich). Cells were incubated consecutively with 30 % Epon in acetone for 1-2 hours, 70 % Epon in acetone for 2 hours and 90 % Epon in acetone o.n at 4°C. The next day, cells were further infiltrated with pure Epon for 8 hours. Epon was changed three times. Later the plastic was polymerized at 60°C for 48 hours. 70 nm sections were cut using a microtome (Reichert Ultracut S, Reichert/Leica, Wetzlar, Germany) and collected on 0.5% formvar grids (G2200C, Plano, Wetzlar Germany). The sections were stained with 2.5% uranyl acetate in 70% methanol for 5 min prior to imaging. Sections were imaged on a TM-Zeiss-900 electron microscope equipped with a 1k slow scan CCD camera (TW 7888, PROSCAN, Germany). Presynaptic structures were identified by morphology. Image analysis was performed using custom macros (ImageJ). Briefly, vesicle and endosome structures were outlined with the “freehand” selection tool and area and length was measured. In addition a line was drawn along the PSD and measured. Docked vesicles were counted manually. Vesicles were defined as docked when there vesicle membrane touched the plasma membrane. Data points were plotted with GraphPad Prism. Statistical significance was determined using t-test with GraphPad Prism.

### Immunocytochemistry (ICC)

Hippocampal neurons DIV14-16 were prepared for ICC. In brief, cells growing on coverslips were washed in PBS and subsequently fixed in 4% paraformaldehyde (PFA) for 5 min at RT. After washing in phosphate buffered saline (PBS, Thermo Fisher scientific, Waltham, USA), cells were permeabilized with 0.1 % Tween20 in PBS (PBS-T). Afterwards cells were blocked in blocking solution (5 % normal goat serum in PBS-T) for 30 min and incubated with primary antibodies in blocking solution overnight at 4°C. The following antibodies were used: Synaptophysin (1:1000; mouse; synaptic systems, Göttingen, germany; Cat# 101011, RRID:AB_887824), Synaptophysin (1:1000; guinea pig; synaptic systems, Göttingen, germany; Cat# 101004, RRID:AB_1210382), Synapsin (1:200; rabbit; abcam, Cambridge, UK; Cat# ab64581, RRID:AB_1281135), VGlut1 (1:1000; guinea pig; synaptic systems, Göttingen, germany; Cat# 135304, RRID:AB_887878), Rab5 (1:500; mouse; synaptic systems, Göttingen, germany; Cat# 108011, RRID:AB_887773), Rab7 (1:500; rabbit; abcam, Cambridge, UK; Cat# ab137029, RRID:AB_2629474), EEA1 (1:200; rabbit; cell signaling, Danvers, USA; Cat# 3288S, RRID:AB_2096811), GFP (1:500; chicken; Thermo Scientific, Waltham, USA; Cat# A10262, RRID:AB_2534023), GFP (1:500; mouse; Roche, Basel, Switzerland; Cat# 11814460001, RRID:AB_390913), Rabex5 (1:100; rabbit; Thermo scientific, Waltham, USA; Cat# PA5-21117, RRID:AB_11157010) MAP2 (1:1000; chicken; Millipore, Darmstadt, Germany; Cat# AB5543, RRID:AB_571049), Piccolo (1:500; rabbit; synaptic systems, Göttingen, germany; Cat# 142002, RRID:AB_887759), Piccolo (1:1000; rabbit; abcam, Cambridge, UK; Cat# ab20664, RRID:AB_777267), Piccolo (1:500, guinea pig, synaptic systems, Göttingen, Germany; Cat# 142104, RRID:AB_2619831). Afterwards cells were washed 3 times with PBS-T and incubated with secondary antibodies for 1 h at RT. Differently labeled secondary antibodies were used from Invitrogen (Thermo Fisher scientific, Waltham, USA, dilution 1:1000). After washing, coverslips were mounted in ProLong Diamond Antifade Mountant (Thermo Fisher scientific, Waltham, USA).

### Super-resolution imaging using structured illumination

SIM imaging was performed on a Deltavision OMX V4 microscope equipped with three watercooled PCO.edge sCMOS cameras, 405 nm, 488 nm, 568 nm and 642 nm laserlines and a ×60 1.42 numerical aperture Plan Apochromat lense (Olympus). z-Stacks covering the whole cell, with sections spaced 0.125 μm apart, were recorded. For each z-section, 15 raw images (three rotations with five phases each) were acquired. Final super-resolution images were reconstructed using softWoRx software and processed in ImageJ/FIJI.

### FM-dye uptake

Functional presynaptic terminals were labeled with FM4-64 dye (Invitrogen, Thermo Fisher scientific, Waltham, USA) as described earlier (Waites et al., 2013). In brief, neurons 12-14 DIV were mounted in a custom-built chamber perfused with Tyrode`s saline solution (25 mM HEPES, 119 mM NaCl, 2.5 mM KCl, 30 mM glucose, 2 mM CaCl_2_, 2 mM MgCl_2_, pH 7.4) at 37°C. An image of the basal background fluorescence was taken before the addition of the dye. To define the total recycling pool of SV, neurons were incubated with Tyrode`s buffer containing 1μg/ml FM4-64 dye and 60 or 90 mM KCl for 90 s. Subsequently unbound dye was washed off and images were acquired from different areas of the coverslip. 10 µM Latrunculin A or 5 µM Jasplakinolide (both Calbiochem) was added to Tyrode’s solution prior to stimulation for 5 min or 10 min, respectively.

### FM-dye destain

DIV 21 *Pclo*^*wt/wt*^ as well as *Pclo*^*gt/gt*^ primary hippocampal neurons were mounted in a field stimulation chamber (Warner Instruments, Hamden, USA) and perfused with Tyrode’s saline solution (25 mM HEPES, 119 mM NaCl, 2.5 mM KCl, 30 mM glucose, 2 mM CaCl2, 2 mM MgCl2, pH 7.4) at 37°C. At first an image was taken to obtain background fluorescence. Afterwards Tyrode’s saline solution was exchanged with Tyrode’s saline solution plus 1μg/ml FM1-43 dye and neurons were stimulated with 900 AP 10 Hz using a field stimulator (Warner Instruments, Hamden, USA) which was controlled by pCLAMP Software (Molecular Devices, Sunnyvale, USA). Subsequent stimulation unbound dye was washed away floating at least 10 ml Tyrode’s saline solution through the chamber. Following washing 5 images were taken to document FM 1-43 dye uptake during stimulation. Afterwards cells were stimulated again with 900 AP and 10Hz. 120 images were taken every second to document unloading of the FM 1-43 dye during stimulation.

### Image acquisition

Images for immunocytochemical stainings as well as FM uptake experiments were acquired on a spinning disc confocal microscope (Zeiss Axio Observer.Z1 with Andor spinning disc unit and cobolt, omricron, i-beam laser (405, 490, 562 and 642 nm wavelength)) using a 63x 1.4 NA Plan-Apochromat oil objective and an iXon ultra (Andor, Belfast, UK) camera controlled by iQ software (Andor, Belfast, UK). For live cell imaging, neurons (DIV14–16) or HEK293 cells growing on 22 × 22 mm coverslips were mounted in a custom-built chamber designed for perfusion, heated to 37°C by forced-air blower and perfused with Tyrode’s saline solution (25 mM HEPES, 119 mM NaCl, 2.5 mM KCl, 30 mM glucose, 2 mM CaCl_2_, 2 mM MgCl_2_, pH 7.4). Images for FM destain experiments were acquired on a Olympus IX83 microscope (Olympus, Hamburg, Germany) equipped with a 60x 1.2 NA UPlanSApo water objective, a CoolLED system (405 nm, 470 nm, 555 nm, 640 nm wavelength) (Acal^bfi^, Göbenzell, Germany) and a Andor Zyla camera (Andor, Belfast, UK).

### Image processing

For image processing ImageJ/FIJI and OpenView software was used (Schindelin et al., 2012). Piccolo intensity in VGlut1 puncta as well as FM4-64 dye uptake was measured using a box routine with OpenView software (written by Dr. Noam Ziv, Technion Institute, Haifa, Israel). To measure Synaptophysin intensity in *Pclo*^*wt/wt*^ and *Pclo*^*gt/gt*^ neurons, 9-pixel regions of interest (ROIs) positive for Synaptophysin were manually selected in ImageJ, subsequently the mean intensity within these ROIs was measured using a customized ImageJ script. The number of Rab5-GFP, Rab7-GFP and EEA1 positive puncta along axons was determined from randomly picked axon sections. Numbers of puncta per unit length were counted manually. The intensity of various endosome proteins (Rabex5, Rab5, EEA1, Pra1) at synapses was measured using a customized ImageJ script. 9-pixel ROIs were manually picked based on Synaptophysin staining. Subsequently corresponding fluorescence intensities were measured in all active channels using a the customized ImageJ script. The background fluorescence was substracted from all ROIs before the Average intensity of the endosome proteins was calculated from all selected ROIs. Negative intensities were counted as 0. The fraction of synapses positive for endosome proteins was calculated with a defined intensity value as threshold. It was defined as the standard derivation intensity of the corresponding endosome protein measured in *Pclo*^*wt/wt*^ synapses. The size of GFP-2x-FYVE positive compartments imaged with SIM was measured using ImageJ. The “freehand” tool was used to mark and measure the area of GFP-2x-FYVE puncta. The intensity of endosome proteins (Rabex5, Rab5, EEA1) at GFP-2x-FYVE puncta was measured using a customized ImageJ script. 9-pixel ROIs were picked manually based on GFP-2x-FYVE staining along axons. Subsequently corresponding fluorescence intensities were measured in all active channels using a customized ImageJ script. The background fluorescence was substracted from all ROIs before the Average intensity of the endosome protein was calculated from all selected eGFP-2x-FYVE puncta. Negative intensities were counted as 0. The fraction of double or triple positive GFP-2x-FYVE vesicles was calculated using a defined intensity value as threshold. It was defined as the standard derivation intensity of the corresponding protein intensity measured in *Pclo*^*wt/wt*^ synapses. Synaptophysin intensity in the presence or absence of Rab5^Q79L^ was determined in ImageJ. 9-pixel ROIs were picked along axons positive or negative for GFP-2x-FYVE. Subsequently Synaptophysin intensity within the defined ROIs was measured using a customized ImageJ script. The intensity of different endosome proteins at the cell soma was measured using ImageJ. The “free-hand” tool was used to surround the soma. Subsequently the fluorescence intensity within this area was measured and depicted as intensity per soma area. The intensity of different endosome proteins along dendrites was measured using ImageJ. 9-pixel ROIs were picked manually along dendrites marked by MAP2. Subsequently corresponding fluorescence intensities were measured using a customized ImageJ script.

Post-processing of automatically measured image data used python and the “pandas” data analysis package (McKinney et al., 2010). Statistical analysis was calculated in GraphPad Prism. Data points were plotted using GraphPad Prism.

### Statistical analysis

All values are shown as means ± SEM. Statistical significance was assessed using Student’s *t* test or ANOVA for comparing multiple samples. For all statistical tests, the 0.05 confidence level was considered statistically significant. In all figures, * denotes P < 0.05, ** denotes P < 0.005, *** denotes P < 0.001 and **** denotes P < 0.0001 in an unpaired Student’s *t* test or ANOVA.

## Acknowledgments

We would like to thank Prof. Eckart D. Gundelfinger for discussion and valuable comments on the manuscript, Anny Kretschmer for technical assistance. The Virus Core facility, Charité Berlin. The Advanced Light Microscopy core facility at the Radium Hospital in Oslo for access to an OMX superresolution microscope.

The authors declare no competing financial interests.

## Author contributions

F. Ackermann conceived the study and designed experiments. C. Bruns performed western blot analysis and immunohistochemical stainings. K.O. Schink performed super resolution microscopy, cloned vectors and wrote scripts for analysis. F. Ackermann, C. Rosenmund and C.C. Garner designed experiments. F.K. Hamra and Z. Izsvák generated *Pclo*^*gt/gt*^ rat. F. Ackermann and C.C. Garner wrote the manuscript.

## References

Abdul-Ghani, M., Gougeon, P. Y., Prosser, D. C., Da-Silva, L. F., & Ngsee, J. K. (2001). PRA isoforms are targeted to distinct membrane compartments. J Biol Chem, 276(9), 6225–6233. doi:10.1074/jbc.M009073200

Ackermann, F., Waites, C. L., & Garner, C. C. (2015). Presynaptic active zones in invertebrates and vertebrates. EMBO Rep, 16(8), 923–938. doi:10.15252/embr.201540434

Ahmed, M. Y., Chioza, B. A., Rajab, A., Schmitz-Abe, K., Al-Khayat, A., Al-Turki, S.,… Mochida, G. H. (2015). Loss of PCLO function underlies pontocerebellar hypoplasia type III. Neurology, 84(17), 1745–1750. doi:10.1212/WNL.0000000000001523

Arancillo, M., Min, S. W., Gerber, S., Munster-Wandowski, A., Wu, Y. J., Herman, M.,… Rosenmund, C. (2013). Titration of Syntaxin1 in mammalian synapses reveals multiple roles in vesicle docking, priming, and release probability. J Neurosci, 33(42), 16698–16714. doi:10.1523/JNEUROSCI.0187-13.2013

Balla, T. (2013). Phosphoinositides: tiny lipids with giant impact on cell regulation. Physiol Rev, 93(3), 1019–1137. doi:10.1152/physrev.00028.2012

Banker, G., & Goslin, K. (1988). Developments in neuronal cell culture. Nature, 336(6195), 185–186. doi:10.1038/336185a0

Breustedt, J., Gundlfinger, A., Varoqueaux, F., Reim, K., Brose, N., & Schmitz, D. (2010). Munc13-2 differentially affects hippocampal synaptic transmission and plasticity. Cereb Cortex, 20(5), 1109–1120. doi:10.1093/cercor/bhp170

Campeau, E., Ruhl, V. E., Rodier, F., Smith, C. L., Rahmberg, B. L., Fuss, J. O.,… Kaufman, P. D. (2009). A versatile viral system for expression and depletion of proteins in mammalian cells. PLoS One, 4(8), e6529. doi:10.1371/journal.pone.0006529

Cases-Langhoff, C., Voss, B., Garner, A. M., Appeltauer, U., Takei, K., Kindler, S., Garner, C. C. (1996). Piccolo, a novel 420 kDa protein associated with the presynaptic cytomatrix. Eur J Cell Biol, 69(3), 214–223.

Cherfils, J., & Zeghouf, M. (2013). Regulation of small GTPases by GEFs, GAPs, and GDIs. Physiol Rev, 93(1), 269–309. doi:10.1152/physrev.00003.2012

Choi, K. H., Higgs, B. W., Wendland, J. R., Song, J., McMahon, F. J., & Webster, M. J. (2011). Gene expression and genetic variation data implicate PCLO in bipolar disorder. Biol Psychiatry, 69(4), 353–359. doi:10.1016/j.biopsych.2010.09.042

Christoforidis, S., McBride, H. M., Burgoyne, R. D., & Zerial, M. (1999). The Rab5 effector EEA1 is a core component of endosome docking. Nature, 397(6720), 621–625. doi:10.1038/17618

Davydova, D., Marini, C., King, C., Klueva, J., Bischof, F., Romorini, S.,… Fejtova, A. (2014). Bassoon specifically controls presynaptic P/Q-type Ca(2+) channels via RIM-binding protein. Neuron, 82(1), 181–194. doi:10.1016/j.neuron.2014.02.012

Dick, O., Hack, I., Altrock, W. D., Garner, C. C., Gundelfinger, E. D., & Brandstatter, J. H. (2001). Localization of the presynaptic cytomatrix protein Piccolo at ribbon and conventional synapses in the rat retina: comparison with Bassoon. J Comp Neurol, 439(2), 224–234.

Fenster, S. D., Chung, W. J., Zhai, R., Cases-Langhoff, C., Voss, B., Garner, A. M.,… Garner, C. C. (2000). Piccolo, a presynaptic zinc finger protein structurally related to bassoon. Neuron, 25(1), 203–214.

Fenster, S. D., & Garner, C. C. (2002). Gene structure and genetic localization of the PCLO gene encoding the presynaptic active zone protein Piccolo. Int J Dev Neurosci, 20(3-5), 161-171.

Fenster, S. D., Kessels, M. M., Qualmann, B., Chung, W. J., Nash, J., Gundelfinger, E. D., & Garner, C. C. (2003). Interactions between Piccolo and the actin/dynamin-binding protein Abp1 link vesicle endocytosis to presynaptic active zones. J Biol Chem, 278(22), 20268–20277.

Gillooly, D. J., Morrow, I. C., Lindsay, M., Gould, R., Bryant, N. J., Gaullier, J. M.,… Stenmark, H. (2000). Localization of phosphatidylinositol 3-phosphate in yeast and mammalian cells. Embo J, 19(17), 4577–4588. doi:10.1093/emboj/19.17.4577

Giniatullina, A., Maroteaux, G., Geerts, C. J., Koopmans, B., Loos, M., Klaassen, R.,… Verhage, M. (2015). Functional characterization of the PCLO p.Ser4814Ala variant associated with major depressive disorder reveals cellular but not behavioral differences. Neuroscience, 300, 518–538. doi:10.1016/j.neuroscience.2015.05.047

Girach, F., Craig, T. J., Rocca, D. L., & Henley, J. M. (2013). RIM1alpha SUMOylation is required for fast synaptic vesicle exocytosis. Cell Rep, 5(5), 1294–1301. doi:10.1016/j.celrep.2013.10.039

Grant, B. D., & Donaldson, J. G. (2009). Pathways and mechanisms of endocytic recycling. Nat Rev Mol Cell Biol, 10(9), 597–608. doi:10.1038/nrm2755

Gundelfinger, E. D., Reissner, C., & Garner, C. C. (2015). Role of Bassoon and Piccolo in Assembly and Molecular Organization of the Active Zone. Front Synaptic Neurosci, 7, 19. doi:10.3389/fnsyn.2015.00019

Haferlach, T., & Schoch, C. (2002). [WHO classification of acute myeloid leukaemia (AML) and the myelodysplastic syndrome (MDS)]. Dtsch Med Wochenschr, 127(9), 447–450. doi:10.1055/s-2002-20422

Hagiwara, A., Fukazawa, Y., Deguchi-Tawarada, M., Ohtsuka, T., & Shigemoto, R. (2005). Differential distribution of release-related proteins in the hippocampal CA3 area as revealed by freeze-fracture replica labeling. J Comp Neurol, 489(2), 195–216. doi:10.1002/cne.20633

Harris, T. W., Hartwieg, E., Horvitz, H. R., & Jorgensen, E. M. (2000). Mutations in synaptojanin disrupt synaptic vesicle recycling. J Cell Biol, 150(3), 589–600.

Hayashi, M., Raimondi, A., O’Toole, E., Paradise, S., Collesi, C., Cremona, O.,… De Camilli, P. (2008). Cell- and stimulus-dependent heterogeneity of synaptic vesicle endocytic recycling mechanisms revealed by studies of dynamin 1-null neurons. Proc Natl Acad Sci U S A, 105(6), 2175–2180. doi:10.1073/pnas.0712171105

Heuser, J. E., & Reese, T. S. (1973). Evidence for recycling of synaptic vesicle membrane during transmitter release at the frog neuromuscular junction. J Cell Biol, 57(2), 315–344.

Hoopmann, P., Punge, A., Barysch, S. V., Westphal, V., Buckers, J., Opazo, F.,… Rizzoli, S. O. (2010). Endosomal sorting of readily releasable synaptic vesicles. Proc Natl Acad Sci U S A, 107(44), 19055–19060. doi:10.1073/pnas.1007037107

Horiuchi, H., Lippe, R., McBride, H. M., Rubino, M., Woodman, P., Stenmark, H.,… Zerial, M. (1997). A novel Rab5 GDP/GTP exchange factor complexed to Rabaptin-5 links nucleotide exchange to effector recruitment and function. Cell, 90(6), 1149–1159.

Hutt, D. M., Da-Silva, L. F., Chang, L. H., Prosser, D. C., & Ngsee, J. K. (2000). PRA1 inhibits the extraction of membrane-bound rab GTPase by GDI1. J Biol Chem, 275(24), 18511–18519. doi:10.1074/jbc.M909309199

Izsvak, Z., Frohlich, J., Grabundzija, I., Shirley, J. R., Powell, H. M., Chapman, K. M., Hamra, F. K. (2010). Generating knockout rats by transposon mutagenesis in spermatogonial stem cells. Nat Methods, 7(6), 443–445. doi:10.1038/nmeth.1461

Juranek, J., Mukherjee, K., Rickmann, M., Martens, H., Calka, J., Sudhof, T. C., & Jahn, R. (2006). Differential expression of active zone proteins in neuromuscular junctions suggests functional diversification. Eur J Neurosci, 24(11), 3043–3052. doi:10.1111/j.1460-9568.2006.05183.x

Kim, S., Ko, J., Shin, H., Lee, J. R., Lim, C., Han, J. H.,… Kim, E. (2003). The GIT family of proteins forms multimers and associates with the presynaptic cytomatrix protein Piccolo. J Biol Chem, 278(8), 6291–6300. doi:10.1074/jbc.M212287200

Komada, M., & Soriano, P. (1999). Hrs, a FYVE finger protein localized to early endosomes, is implicated in vesicular traffic and required for ventral folding morphogenesis. Genes Dev, 13(11), 1475–1485.

Leal-Ortiz, S., Waites, C. L., Terry-Lorenzo, R., Zamorano, P., Gundelfinger, E. D., & Garner, C. C. (2008). Piccolo modulation of Synapsin1a dynamics regulates synaptic vesicle exocytosis. J Cell Biol, 181(5), 831–846. doi:10.1083/jcb.200711167

Limbach, C., Laue, M. M., Wang, X., Hu, B., Thiede, N., Hultqvist, G., & Kilimann, M. W. (2011). Molecular in situ topology of Aczonin/Piccolo and associated proteins at the mammalian neurotransmitter release site. Proc Natl Acad Sci U S A, 108(31), E392–401. doi:10.1073/pnas.1101707108

Lippe, R., Miaczynska, M., Rybin, V., Runge, A., & Zerial, M. (2001). Functional synergy between Rab5 effector Rabaptin-5 and exchange factor Rabex-5 when physically associated in a complex. Mol Biol Cell, 12(7), 2219–2228.

Mani, M., Lee, S. Y., Lucast, L., Cremona, O., Di Paolo, G., De Camilli, P., & Ryan, T. A. (2007). The dual phosphatase activity of synaptojanin1 is required for both efficient synaptic vesicle endocytosis and reavailability at nerve terminals. Neuron, 56(6), 1004–1018. doi:10.1016/j.neuron.2007.10.032

Maricich, S. M., Aqeeb, K. A., Moayedi, Y., Mathes, E. L., Patel, M. S., Chitayat, D.,… Zoghbi, H. Y. (2011). Pontocerebellar hypoplasia: review of classification and genetics, and exclusion of several genes known to be important for cerebellar development. J Child Neurol, 26(3), 288–294. doi:10.1177/0883073810380047

Mattera, R., Tsai, Y. C., Weissman, A. M., & Bonifacino, J. S. (2006). The Rab5 guanine nucleotide exchange factor Rabex-5 binds ubiquitin (Ub) and functions as a Ub ligase through an atypical Ub-interacting motif and a zinc finger domain. J Biol Chem, 281(10), 6874–6883. doi:10.1074/jbc.M509939200

McLauchlan, H., Newell, J., Morrice, N., Osborne, A., West, M., & Smythe, E. (1998). A novel role for Rab5-GDI in ligand sequestration into clathrin-coated pits. Curr Biol, 8(1), 34–45.

Meberg, P. J., & Miller, M. W. (2003). Culturing hippocampal and cortical neurons. Methods Cell Biol, 71, 111-127.

Medrano, G. A., Singh, M., Plautz, E. J., Good, L. B., Chapman, K. M., Chaudhary, J.,… Hamra, F. K. (2018). Mutant screen for rat reproduction genes reals Piccolo’s central control over social acuity and brain gonad crosstalk. Nature, submitted.

Milosevic, I., Giovedi, S., Lou, X., Raimondi, A., Collesi, C., Shen, H.,… De Camilli, P. (2011). Recruitment of endophilin to clathrin-coated pit necks is required for efficient vesicle uncoating after fission. Neuron, 72(4), 587–601. doi:10.1016/j.neuron.2011.08.029

Minelli, A., Scassellati, C., Cloninger, C. R., Tessari, E., Bortolomasi, M., Bonvicini, C.,… Gennarelli, M. (2012). PCLO gene: its role in vulnerability to major depressive disorder. J Affect Disord, 139(3), 250–255. doi:10.1016/j.jad.2012.01.028

Minerbi, A., Kahana, R., Goldfeld, L., Kaufman, M., Marom, S., & Ziv, N. E. (2009). Long-term relationships between synaptic tenacity, synaptic remodeling, and network activity. PLoS Biol, 7(6), e1000136. doi:10.1371/journal.pbio.1000136

Mishra, A., Eathiraj, S., Corvera, S., & Lambright, D. G. (2010). Structural basis for Rab GTPase recognition and endosome tethering by the C2H2 zinc finger of Early Endosomal Autoantigen 1 (EEA1). Proc Natl Acad Sci U S A, 107(24), 10866–10871. doi:10.1073/pnas.1000843107

Mukherjee, K., Yang, X., Gerber, S. H., Kwon, H. B., Ho, A., Castillo, P. E.,… Sudhof, T. C. (2010). Piccolo and bassoon maintain synaptic vesicle clustering without directly participating in vesicle exocytosis. Proc Natl Acad Sci U S A, 107(14), 6504–6509. doi:1002307107 [pii]10.1073/pnas.1002307107

Murray, D. H., Jahnel, M., Lauer, J., Avellaneda, M. J., Brouilly, N., Cezanne, A.,… Zerial, M. (2016). An endosomal tether undergoes an entropic collapse to bring vesicles together. Nature, 537(7618), 107–111. doi:10.1038/nature19326

Nishimune, H. (2012a). Active zones of mammalian neuromuscular junctions: formation, density, and aging. Ann N Y Acad Sci, 1274, 24-32. doi:10.1111/j.1749-6632.2012.06836.x

Nishimune, H. (2012b). Molecular mechanism of active zone organization at vertebrate neuromuscular junctions. Mol Neurobiol, 45(1), 1–16. doi:10.1007/s12035-011-8216-y

Nishimune, H., Badawi, Y., Mori, S., & Shigemoto, K. (2016). Dual-color STED microscopy reveals a sandwich structure of Bassoon and Piccolo in active zones of adult and aged mice. Sci Rep, 6, 27935. doi:10.1038/srep27935

Okerlund, N. D., Schneider, K., Leal-Ortiz, S., Montenegro-Venegas, C., Kim, S. A., Garner, L. C.,… Garner, C. C. (2017). Bassoon Controls Presynaptic Autophagy through Atg5. Neuron, 93(4), 897–913 e897. doi:10.1016/j.neuron.2017.01.026

Pavlos, N. J., & Jahn, R. (2011). Distinct yet overlapping roles of Rab GTPases on synaptic vesicles. Small Gtpases, 2(2), 77–81. doi:10.4161/sgtp.2.2.15201

Pfeffer, S., & Aivazian, D. (2004). Targeting Rab GTPases to distinct membrane compartments. Nat Rev Mol Cell Biol, 5(11), 886–896. doi:10.1038/nrm1500

Regus-Leidig, H., Fuchs, M., Lohner, M., Leist, S. R., Leal-Ortiz, S., Chiodo, V. A.,… Brandstatter, J. H. (2014). In vivo knockdown of Piccolino disrupts presynaptic ribbon morphology in mouse photoreceptor synapses. Front Cell Neurosci, 8, 259. doi:10.3389/fncel.2014.00259

Rizzoli, S. O., & Betz, W. J. (2002). Effects of 2-(4-morpholinyl)-8-phenyl-4H-1-benzopyran-4-one on synaptic vesicle cycling at the frog neuromuscular junction. J Neurosci, 22(24), 10680–10689.

Rudnik-Schoneborn, S., Barth, P. G., & Zerres, K. (2014). Pontocerebellar hypoplasia. Am J Med Genet C Semin Med Genet, 166C(2), 173–183. doi:10.1002/ajmg.c.31403

Saheki, Y., & De Camilli, P. (2012). Synaptic vesicle endocytosis. Cold Spring Harb Perspect Biol, 4(9), a005645. doi:10.1101/cshperspect.a005645

Sann, S. B., Crane, M. M., Lu, H., & Jin, Y. (2012). Rabx-5 regulates RAB-5 early endosomal compartments and synaptic vesicles in C. elegans. PLoS One, 7(6), e37930. doi:10.1371/journal.pone.0037930

Sasidharan, N., Sumakovic, M., Hannemann, M., Hegermann, J., Liewald, J. F., Olendrowitz, C.,… Eimer, S. (2012). RAB-5 and RAB-10 cooperate to regulate neuropeptide release in Caenorhabditis elegans. Proc Natl Acad Sci U S A, 109(46), 18944–18949. doi:10.1073/pnas.1203306109

Schindelin, J., Arganda-Carreras, I., Frise, E., Kaynig, V., Longair, M., Pietzsch, T.,… Cardona, A. (2012). Fiji: an open-source platform for biological-image analysis. Nat Methods, 9(7), 676–682. doi:10.1038/nmeth.2019

Schink, K. O., Raiborg, C., & Stenmark, H. (2013). Phosphatidylinositol 3-phosphate, a lipid that regulates membrane dynamics, protein sorting and cell signalling. Bioessays, 35(10), 900–912. doi:10.1002/bies.201300064

Schink, K. O., Tan, K. W., & Stenmark, H. (2016). Phosphoinositides in Control of Membrane Dynamics. Annu Rev Cell Dev Biol. doi:10.1146/annurev-cellbio-111315-125349

Schoch, S., Castillo, P. E., Jo, T., Mukherjee, K., Geppert, M., Wang, Y.,… Sudhof, T. C. (2002). RIM1alpha forms a protein scaffold for regulating neurotransmitter release at the active zone. Nature, 415(6869), 321–326.

Schuske, K. R., Richmond, J. E., Matthies, D. S., Davis, W. S., Runz, S., Rube, D. A.,… Jorgensen, E. M. (2003). Endophilin is required for synaptic vesicle endocytosis by localizing synaptojanin. Neuron, 40(4), 749–762.

Siksou, L., Rostaing, P., Lechaire, J. P., Boudier, T., Ohtsuka, T., Fejtova, A.,… Marty, S. (2007). Three-dimensional architecture of presynaptic terminal cytomatrix. J Neurosci, 27(26), 6868–6877.

Simonsen, A., Lippe, R., Christoforidis, S., Gaullier, J. M., Brech, A., Callaghan, J.,… Stenmark, H. (1998). EEA1 links PI(3)K function to Rab5 regulation of endosome fusion. Nature, 394(6692), 494–498. doi:10.1038/28879

Smith, C. B., & Betz, W. J. (1996). Simultaneous independent measurement of endocytosis and exocytosis. Nature, 380(6574), 531–534. doi:10.1038/380531a0

Spang, A. (2009). On the fate of early endosomes. Biol Chem, 390(8), 753–759. doi:10.1515/BC.2009.056

Stenmark, H. (2009). Rab GTPases as coordinators of vesicle traffic. Nat Rev Mol Cell Biol, 10(8), 513–525. doi:10.1038/nrm2728

Stenmark, H., Parton, R. G., Steele-Mortimer, O., Lutcke, A., Gruenberg, J., & Zerial, M. (1994). Inhibition of rab5 GTPase activity stimulates membrane fusion in endocytosis. Embo J, 13(6), 1287–1296.

Stenmark, H., Vitale, G., Ullrich, O., & Zerial, M. (1995). Rabaptin-5 is a direct effector of the small GTPase Rab5 in endocytic membrane fusion. Cell, 83(3), 423–432.

Sudhof, T. C., & Rizo, J. (2011). Synaptic vesicle exocytosis. Cold Spring Harb Perspect Biol, 3(12). doi:10.1101/cshperspect.a005637

Sullivan, P. F., de Geus, E. J., Willemsen, G., James, M. R., Smit, J. H., Zandbelt, T.,… Penninx, B. W. (2009). Genome-wide association for major depressive disorder: a possible role for the presynaptic protein piccolo. Mol Psychiatry, 14(4), 359–375. doi:mp2008125 [pii] 10.1038/mp.2008.125

Takao-Rikitsu, E., Mochida, S., Inoue, E., Deguchi-Tawarada, M., Inoue, M., Ohtsuka, T., & Takai, Y. (2004). Physical and functional interaction of the active zone proteins, CAST, RIM1, and Bassoon, in neurotransmitter release. J Cell Biol, 164(2), 301–311.

Tanaka, H. (2002). [Culturing hippocampal neurons]. Nihon Yakurigaku Zasshi, 119(3), 163–166.

Terry-Lorenzo, R. T., Torres, V. I., Wagh, D., Galaz, J., Swanson, S. K., Florens, L.,… Garner, C. C. (2016). Trio, a Rho Family GEF, Interacts with the Presynaptic Active Zone Proteins Piccolo and Bassoon. PLoS ONE, 11(12), e0167535. doi:10.1371/journal.pone.0167535

Uytterhoeven, V., Kuenen, S., Kasprowicz, J., Miskiewicz, K., & Verstreken, P. (2011). Loss of skywalker reveals synaptic endosomes as sorting stations for synaptic vesicle proteins. Cell, 145(1), 117–132. doi:10.1016/j.cell.2011.02.039

Verstreken, P., Koh, T. W., Schulze, K. L., Zhai, R. G., Hiesinger, P. R., Zhou, Y.,… Bellen, H. J. (2003). Synaptojanin is recruited by endophilin to promote synaptic vesicle uncoating. Neuron, 40(4), 733–748.

Wagh, D., Terry-Lorenzo, R., Waites, C. L., Leal-Ortiz, S. A., Maas, C., Reimer, R. J., & Garner, C. C. (2015). Piccolo Directs Activity Dependent F-Actin Assembly from Presynaptic Active Zones via Daam1. PLoS ONE, 10(4), e0120093. doi:10.1371/journal.pone.0120093

Waites, C. L., Leal-Ortiz, S. A., Andlauer, T. F., Sigrist, S. J., & Garner, C. C. (2011). Piccolo regulates the dynamic assembly of presynaptic F-actin. J Neurosci, 31(40), 14250–14263. doi:31/40/14250 [pii] 10.1523/JNEUROSCI.1835-11.2011

Waites, C. L., Leal-Ortiz, S. A., Okerlund, N., Dalke, H., Fejtova, A., Altrock, W. D.,… Garner, C. C. (2013). Bassoon and Piccolo maintain synapse integrity by regulating protein ubiquitination and degradation. Embo J, 32(7), 954–969. doi:10.1038/emboj.2013.27

Wang, X., Hu, B., Zieba, A., Neumann, N. G., Kasper-Sonnenberg, M., Honsbein, A.,… Kilimann, M. W. (2009). A protein interaction node at the neurotransmitter release site: domains of Aczonin/Piccolo, Bassoon, CAST, and rim converge on the N-terminal domain of Munc13-1. J Neurosci, 29(40), 12584–12596. doi:29/40/12584 [pii] 10.1523/JNEUROSCI.1255-09.2009

Wang, X., Kibschull, M., Laue, M. M., Lichte, B., Petrasch-Parwez, E., & Kilimann, M. W. (1999). Aczonin, a 550-kD putative scaffolding protein of presynaptic active zones, shares homology regions with Rim and Bassoon and binds profilin. J Cell Biol, 147(1), 151–162.

Watanabe, S., Rost, B. R., Camacho-Perez, M., Davis, M. W., Sohl-Kielczynski, B., Rosenmund, C., & Jorgensen, E. M. (2013). Ultrafast endocytosis at mouse hippocampal synapses. Nature, 504(7479), 242–247. doi:10.1038/nature12809

Woudstra, S., van Tol, M. J., Bochdanovits, Z., van der Wee, N. J., Zitman, F. G., van Buchem, M. A.,… Hoogendijk, W. J. (2013). Modulatory effects of the piccolo genotype on emotional memory in health and depression. PLoS One, 8(4), e61494. doi:10.1371/journal.pone.0061494

Wucherpfennig, T., Wilsch-Brauninger, M., & Gonzalez-Gaitan, M. (2003). Role of Drosophila Rab5 during endosomal trafficking at the synapse and evoked neurotransmitter release. J Cell Biol, 161(3), 609–624. doi:10.1083/jcb.200211087

Zerial, M., & McBride, H. (2001). Rab proteins as membrane organizers. Nat Rev Mol Cell Biol, 2(2), 107–117. doi:10.1038/35052055

